# Changes in habitat connectivity for range-restricted birds reflect patterns of woodland invasion

**DOI:** 10.1101/2025.07.18.665495

**Authors:** Arunima Jain, Chiti Arvind, V. Jobin, Abhimanyu Lele, V.V. Robin

## Abstract

Habitat fragmentation and landscape change are common causes of concern for species persistence, especially for habitat specialists. The composition of the matrix surrounding habitat fragments influences connectivity between them, which affects gene flow across the landscape. This can further impact populations in various ways. The Shola Sky Island landscapes naturally comprise a biphasic, forest-grassland mosaic ecosystem unique to the high-altitude regions of the Western Ghats of India. Earlier, this mosaic consisted of patches of native cloud forests embedded in a large grassland matrix. Over the last few decades, however, extensive invasion (up to 60%) of timber species into native grasslands has inverted this mosaic, i.e., small patches of grasslands are nested in a woodland matrix. We attempt to study the effects of these modifications on functional habitat connectivity in this region by modelling species movement using a circuit theory-based algorithm. We do this for seven Shola endemic, range-restricted bird species; six forest-specialist and one grassland-specialist species, based on a decade of field data. We consider a range of species-environment relationships and dispersal capacities for a past, relatively uninvaded landscape and a present, highly modified landscape. We used bird occupancy data (presence/absence from a total of 720 grid cells from targeted occupancy surveys for forest species and 744 presence locations for grassland species from occupancy surveys combined with opportunistic records) along with remotely sensed landscape, vegetation, climatic and topographic variables. We find that connectivity has increased overall for forest specialists, but has reduced for the grassland species. This pattern is concordant with regions where woodland cover from invasive timber species has expanded over approximately two decades. We also identify species-specific areas of low and high connectivity, which may have implications for gene flow within the landscape. This would help focus conservation efforts and predict how future landscape change might affect species persistence.

## Introduction

Natural landscapes and ecosystems are changing globally due to human-led landscape change, which poses a significant threat to the persistence of many animal species (Newbold et al., 2015). The increasing fragmentation and destruction of natural habitats due to anthropogenic landscape change causes populations to become more isolated, which could lead to reduced gene flow between populations. This, in turn, leads to diminishing population sizes due to reduced genetic diversity (MacDougall-Shackleton et al., 2011; Templeton et al., 1990) and possibly local extinctions (reviewed in Turner, 1996). However, not all changes are negative; certain species may be less affected (Blanchet et al., 2010) or even respond positively to landscape change (Carrara et al., 2015; Coppedge et al., 2001). These responses vary based on species ecology. For example, habitat-specialist species are shown to be especially affected by landscape change (Devictor et al., 2008), owing to their narrow niche breadths. Thus, there is a growing need to identify wildlife corridors crucial for movement across a range of species ecologies and protect regions of high movement, to preserve connectivity between populations across their range of distribution (Haddad et al., 2015; Krosby et al., 2010).

The effect of anthropogenic change on habitat connectivity can also be amplified in regions of limited spatial scales, such as montane “sky island” systems, where valleys act as geographic and climatic barriers, isolating mountaintops from each other and from lower elevation regions (Love et al., 2023; Robin, Vishnudas, et al., 2015). The Shola Sky Islands of the Western Ghats of India are one such example. This landscape has fostered the evolution of a unique assemblage of species, endemic to each of these high-elevation mountains of the Western Ghats (Myers et al., 2000; Robin, Vishnudas, et al., 2015; Robin & Nandini, 2012). These range-restricted species may be especially vulnerable to landscape and climate change (Dirnböck et al., 2011).

Over the last few decades, the Shola Sky Island landscapes have undergone extensive modifications. Historically, they consisted of a biphasic mosaic habitat, where fragments of native Shola forest were surrounded by a large native montane grassland (Robin & Nandini, 2012), in the form of alternate stable states (Joshi et al., 2020). However, with these changes, this mosaic has been highly modified. Native grasslands have particularly been impacted, with the expansion of human settlements, and the subsequent plantation of timber species that turned invasive, such as *Acacia*, *Eucalyptus* and *Pinus* species, along with the establishment of a production landscape (tea, coffee, cardamom, etc.) (Arasumani et al., 2018, 2019; Joshi et al., 2018). This has subsequently affected the distributions of several endemic species across the landscape (Jobin et al., 2025; Lele et al., 2020, 2024; Sukumar et al., 1995).

Landscape change in a habitat mosaic can impact species movement and populations in several ways. It can alter the composition and configuration of the matrix surrounding suitable habitat patches, making it more or less suitable for movement and/or occupancy. This could influence individual dispersal and population composition in different ways (reviewed in Fletcher et al., 2024). It can also alter the spatial configuration of the habitat mosaic itself, through changes in the total area of suitable habitat, or through changes in the fragmentation levels of suitable habitat, which are most often highly correlated (Cushman et al., 2012). The unique stable biphasic nature of the Shola Sky Island landscape has allowed different kinds of habitat specialists and generalists to coexist in close proximity to each other (Robin & Nandini, 2012). We may thus see landscape change influencing movement, and subsequently population connectivity, in different ways within the same landscape and climate space, i.e., through changing the matrix composition, as well as the configuration of the habitat mosaic.

Anthropogenic activities in this region have not only led to an inversion of the forest-grassland mosaic, where grassland patches are now nested within a woodland matrix, but have also introduced several new land use types into the landscape (Arasumani et al., 2019). This has led to the creation of a more heterogeneous landscape, which may be perceived differently by different species. The Shola Sky Island landscape, therefore, provides an interesting perspective in terms of how habitat connectivity for different species may be affected by changes in an already naturally fragmented mosaic habitat (Robin, Gupta, et al., 2015).

In this study, we aim to estimate the extent of change in habitat connectivity for forest and grassland specialist bird species across two Sky Islands, the Nilgiris and the Palani-Anamalai hills. These are the two largest Sky Islands of the Western Ghats and have been major sites for human settlement and human-led landscape change in the Western Ghats over the last approximately 200 years in the Nilgiris (Joshi et al., 2018) and 50 years in the Palani-Anamalai region (Arasumani et al., 2018, 2019; Mudappa & Raman, 01 2007). We specifically ask a) if changes in habitat connectivity corroborate land cover change due to human-led modifications and invasion, and b) identify species-specific regions of change with implications for conservation action.

We expect that overall connectivity across the landscape may have increased for the forest specialists and decreased for the grassland specialist due to the encroachment of open grasslands by woody invasive species, closely following patterns of niche use for these species (Jobin et al., 2025; Lele et al., 2020). We expect these changes to have concentrated in regions of high land use change over the last few decades.

## Methods

### Study landscape and occupancy data

Our study landscape covers approximately 2894 sq. km. across the two largest Sky Islands in the southern Western Ghats above 1400m ASL. This includes the Nilgiris and the Palani-Anamalais, situated north and south of the Palghat Gap, respectively, a valley which acts as a major biogeographic barrier to multiple species (Robin, Vishnudas, et al., 2015; Sekar & Karanth, 2013). Presently, both these landscapes, particularly the eastern parts, are highly anthropogenically modified (Arasumani et al., 2019). Most of the western half of the Nilgiris and the Anamalai region of the Palani-Anamalais are designated as Protected Areas and have remained intact (Arasumani et al., 2019). There is also a climatic gradient along a longitudinal axis (Caner et al., 2007; Pascal, 1982; Sekar & Karanth, 2013), with a shift in precipitation seasonality, going from the west to the east.

The landscape modifications in the Nilgiris are much older, owing to the afforestation of grasslands and the establishment of a tea production landscape more than 150 years ago (Joshi et al., 2018; Ramesh et al., 2025). Landscape change in the Palani-Anamalai region, however, starting from the 1970s, has resulted in over 60% of the landscape being modified, owing to the rapid expansion of timber plantations that have turned invasive and further taken over the native grasslands (Arasumani et al., 2019). These non-native woodlands have facilitated the invasion of several other non-native plants in their understory (Jobin et al., 2023), and thus have a different composition from native Shola forests. Some older timber plantations also allow the regeneration of native Shola forest species in their understory (Schmerbeck et al., 2024), further facilitating the expansion of wooded habitats.

Occupancy data for forest species were obtained from field surveys conducted by Jobin et al. (2025). Of the 10 bird species targeted in these surveys, we selected six forest specialist species for this study since their global distributions are strictly limited to the high-elevation regions in the southern Western Ghats – the Shola Sky Islands, making them potentially more vulnerable to landscape and climate change (Dirnböck et al., 2011). These six species are the White-bellied Sholakili (*Sholicola albiventris*, henceforth SHAL) and Palani Laughingthrush (*Montecincla fairbanki*, henceforth MOFA), found only in the Palani-Anamalai and Highwavies region; the Nilgiri Sholakili (*Sholicola major*, henceforth SHMA), found mostly in the Nilgiris region and the Nilgiri Laughingthrush (*Montecincla cachinnans*, henceforth MOCA), found only in the Nilgiris region; and the Black-and-orange Flycatcher (*Ficedula nigrorufa*, henceforth FINI) and Nilgiri Flycatcher (*Eumyias albicaudatus*, henceforth EUAL), found in both regions. We used presence/absence data collected from 397 grid cells in the Palani-Anamalai region and 323 grid cells in the Nilgiris at a 100m x 100m scale, in regions higher than 1400m ASL, to model habitat suitability for these species.

For the grassland specialist species Nilgiri Pipit (*Anthus nilghiriensis*, henceforth ANNI), we used a composite dataset of presence points recorded during surveys as well as opportunistic records by Lele et al. (2020, 2024). We filtered these points to one point per grid cell to avoid sampling bias (Boria et al., 2014), using the *biomod2* package in R (Thuiller et al., 2025), to get 744 points. We then randomly generated 5000 pseudo-absences from the background to represent most of the potentially unused niches, such that they were at least 1000m away from any presence point using the *biomod2* package, to avoid similar niches to those of presences (Barbet-Massin et al., 2012). Refer to Table S2 in the Supplementary Material for more information about occupancy data for all species and (Jobin et al., 2025; Lele et al., 2020, 2024) for detailed information on data collection methods.

### Source strength data

Our study area covered a large extent, encompassing most of the global distribution of all seven species. Thus, it was not feasible to collect data on actual abundances across our entire region of interest. Instead, we used habitat suitability as a proxy for source strength in the connectivity model. We extracted Shola forest patches for forest species and native grassland patches for the grassland species from habitat suitability layers generated using niche modelling methods (described below), to use as source strength layers for the connectivity models.

To generate habitat suitability layers, we used predictions of the probability of presence of a species from Random Forest models (Breiman, 2001), implemented using the *randomForest* package in R (Liaw & Wiener, 2002). We constructed these models using occupancy data collected from field surveys as the response variable and remotely sensed environmental variables as predictors. Random Forest is one of the most commonly used machine learning algorithms in landscape ecology (Stupariu et al., 2022) and has been shown to generally outperform most other algorithms used to construct environmental niche models, especially traditional regression-based models (Elith et al., 2020; Valavi et al., 2022; Wunderlich et al., 2022). It can handle non-parametric and complex, non-linear relationships as well as variable interactions, often necessary when modelling species-environment relationships (Cutler et al., 2007; Evans & Cushman, 2009), with minimal model tuning required (Freeman et al., 2016; Probst et al., 2018).

For the grassland specialist species, we used a down-sampled Random Forest model to predict the probability of presence, i.e., the number of pseudo-absences randomly sampled from the training dataset in each bootstrap sample was kept equal to the number of presences.

Down-sampling the majority class has been shown to improve the prediction accuracy of Random Forest models and helps to avoid model overfitting, especially when using presence-only data in species distribution models (Evans & Cushman, 2009; Robinson et al., 2018; Valavi et al., 2022).

We resampled predictor variables for all species to a 100m x 100m resolution to match the targeted sampling grid size of the forest species. We removed variables that were correlated with others (Pearson’s correlation coefficient > 0.7) before training each model. Refer to Table S1 for a list of all used variables, with their descriptions and sources, and Table S2 for model performance scores and tuning parameters, in the Supplementary Material.

### Resistance layers

We generated resistance layers for the connectivity model by taking the inverse of habitat suitability layers using the following transformation formula from Trainor et al. (2013):

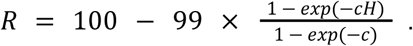

This method has been proven to accurately represent landscape resistance to movement in the absence of field data on species movement (Ahmadi et al., 2017; Keeley et al., 2016; Valerio et al., 2019), and especially for specialist species (Keeley et al., 2016; Trainor et al., 2013). Here, *R* represents resistance to movement on a scale of 1 to 100, *H* represents habitat suitability, and *c* is a constant that determines the non-linearity of the transformation function. A *c* value of 0.25 generates a close-to-linear negative transformation, and *c* > 0.25 generates negative exponential transformations, getting steeper as you increase *c* (Trainor et al., 2013). Most species are likely to move across otherwise unsuitable or moderately suitable habitats during dispersal. Thus, a negative exponential transformation of habitat suitability generally represents resistance to movement more accurately than a linear transformation (Keeley et al., 2016, 2017; Trainor et al., 2013). We generated three resistance layers for each species, with *c* values of 0.25, 2 and 8, respectively, to model connectivity across a range of potential dispersal scenarios going from low to high flexibility with respect to habitat selection during dispersal.

### Past projection

We assume species have not changed in their ecology significantly between the recent past and the present, and track the same environmental variables in the same ways. Therefore, to compare habitat connectivity in a present, highly modified landscape with a past, relatively unmodified landscape, we projected current predictions of the probability of presence to the oldest time point for which data on the same environmental variables were available. Habitat suitability layers and the corresponding source strength and resistance layers were generated for both Sky Island landscapes at a time point between 20 to 25 years before the current period of study (2019 - 2023). For these projections, predictor variables derived from the 2017 land use dataset were replaced with the same predictor variables derived from the 1995 land use dataset (Arasumani et al., 2019), and canopy height data from the year 2020 were replaced by canopy height data from the year 2000 (Potapov et al., 2021). Refer to Table S1 in the supplementary material for more information about the variables.

### Connectivity models

We used a circuit theory-based algorithm, Omniscape (Landau et al., 2021), implemented in Julia 1.10.0 (Bezanson et al., 2017), to model species movement in the manner of electrical current flowing across a resistance surface. It is a generalised version of the Circuitscape algorithm (Hall et al., 2021), which models movement between pre-defined pairs of cores of suitable habitat. Omniscape uses a more flexible approach and implements the algorithm iteratively over a moving window of a fixed radius around each source pixel (Landau et al., 2021). Thus, it is more useful for modelling habitat connectivity where suitable habitats may not be defined as discrete patches, i.e. habitat gradients are more continuous (Landau et al., 2021; B. McRae et al., 2016), while accounting for the dispersal capacity of the target species.

We constructed a total of 126 models to model movement for seven species (6 forest and 1 grassland specialist) for 9 dispersal scenarios (3 resistance layers x 3 moving window radii) in both a present-day, highly modified landscape and a past, relatively unmodified landscape. To set the radius of the moving window based on dispersal capacity for each species, we used knowledge from field experts, preliminary data from our long-term monitoring sites in the field and literature on the closest related bird species or other bird species with similar habitat requirements and ecologies (Ceresa et al., 2023; Ehlers Smith et al., 2018; Fagan et al., 2016; Şekercioğlu et al., 2015). We arrived at the following ranges of radii for each species: 1000m, 3000m and 5000m for ANNI; 500m, 1000m, and 1500m for EUAL; and 200m, 500m, and 1000m for the rest.

Omniscape produces three output layers, in the form of cumulative current flow (CCF), flow potential (FP), and normalised cumulative current flow (NCF) maps. CCF reflects the relative number of potential paths that pass through each pixel, acting as a proxy for movement suitability (McRae et al., 2008). FP represents current flow (movement) in a null model scenario, when the entire region is uniformly resistant. NCF is derived from CCF divided by FP, and thus represents current flow relative to a null model scenario. Thus, regions with values of NCF > 1 represent channelised current flow or potential movement corridors and pinch points, NCF < 1 represent impeded current flow or potential barriers to movement, and NCF ≈ 1 represent diffuse current flow, i.e., movement is neither impeded nor channelised (Landau et al., 2021; B. McRae et al., 2016). We plotted NCF maps for each model, after binning these values into percentiles, to visualise habitat connectivity in all models, by mapping potential barriers and corridors for movement. We then subtracted past CCF values from present CCF values for each model to visualise changes in movement suitability over time.

To compare and study spatial patterns of change in connectivity and present connectivity across species, we divided each Sky Island into 50 longitudinal slices of equal width and summarised change in CCF and present-day CCF for each slice by taking the median. We then plotted these values after smoothing using a Generalised Additive Model (GAM) along the east-west axis using the *ggplot2* package in R (Wickham, 2016). We also combined the 2017 and 1995 land use maps from Arasumani et al. (2019) to classify landscape change into 8 categories (Historical human-use and water bodies, Woodland to human-use, Grassland to human-use, Grassland gain, Grassland to woodland, Human-use to woodland, Stable grassland and Stable woodland) to compare with patterns of change in connectivity. Here, “human-use” land cover categories represent tea plantations, other agricultural land, and settlements, and “woodland” represents both native Shola and non-native woodland cover.

## Results

### Spatial patterns of present connectivity

We visualised movement patterns for 7 species using the Omniscape algorithm from occupancy data from 720 grid cells for 6 forest specialist species and 744 presence records for one grassland specialist species across two of the largest Sky Islands in the Western Ghats of India, along with remotely sensed environmental data. We identified potential barriers and pinch points for movement across both landscapes using normalised current flow (NCF) maps (Fig. 2). We found that movement seems to be diffuse across most of the landscape, except for around the edges, which may be attributed to the surrounding missing cells being set to the highest resistance.

**Fig. 1:**
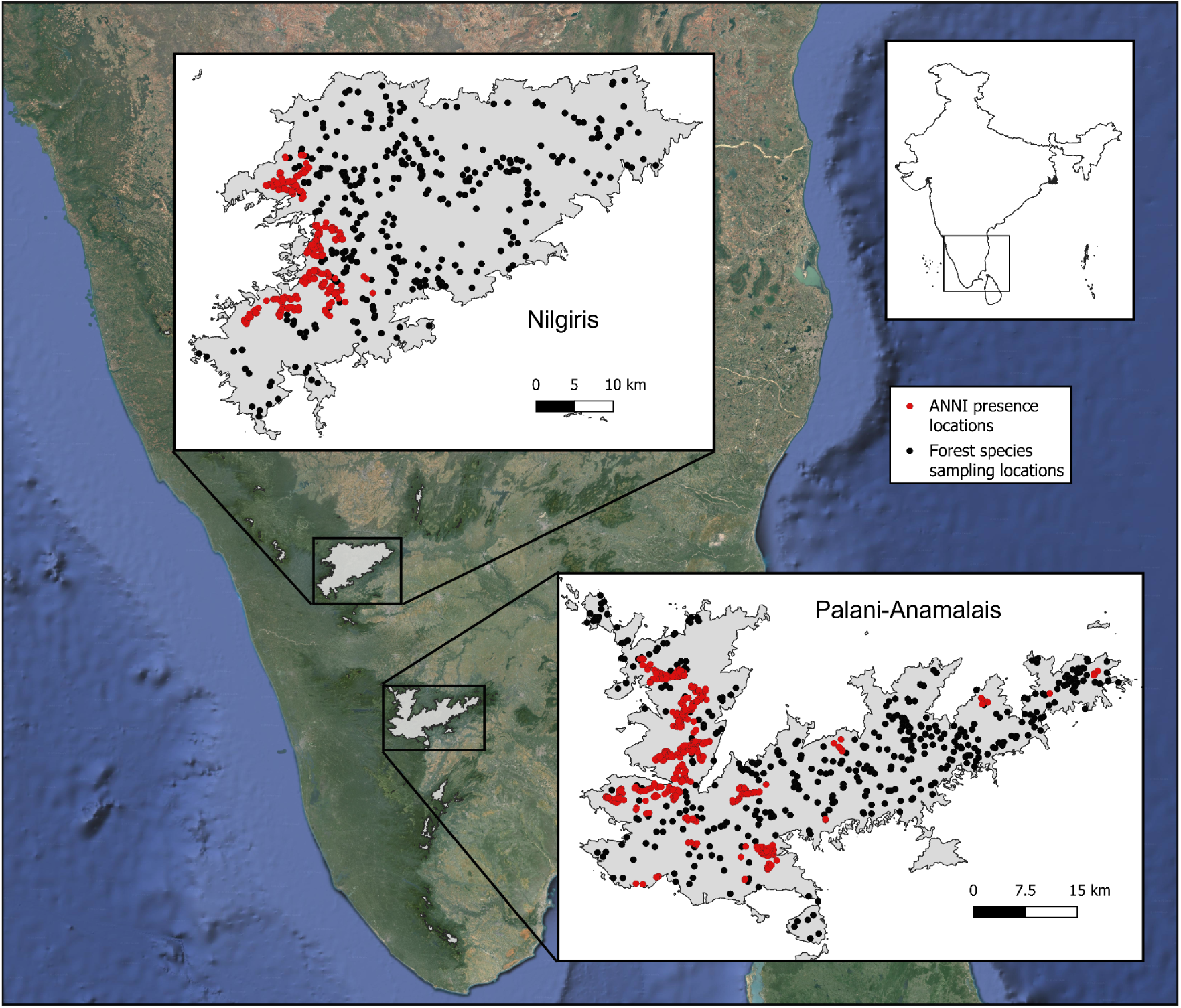
Study landscape and occupancy data. Black dots represent sampling locations for all forest species: 397 grid cells in the Palani-Anamalai region and 323 grid cells in the Nilgiri region. Red dots represent 744 presence locations across both regions of the grassland species Nilgiri Pipit (*Anthus nilghiriensis*). Grey polygons represent regions of the Western Ghats higher than 1400m ASL.

**Fig. 2:**
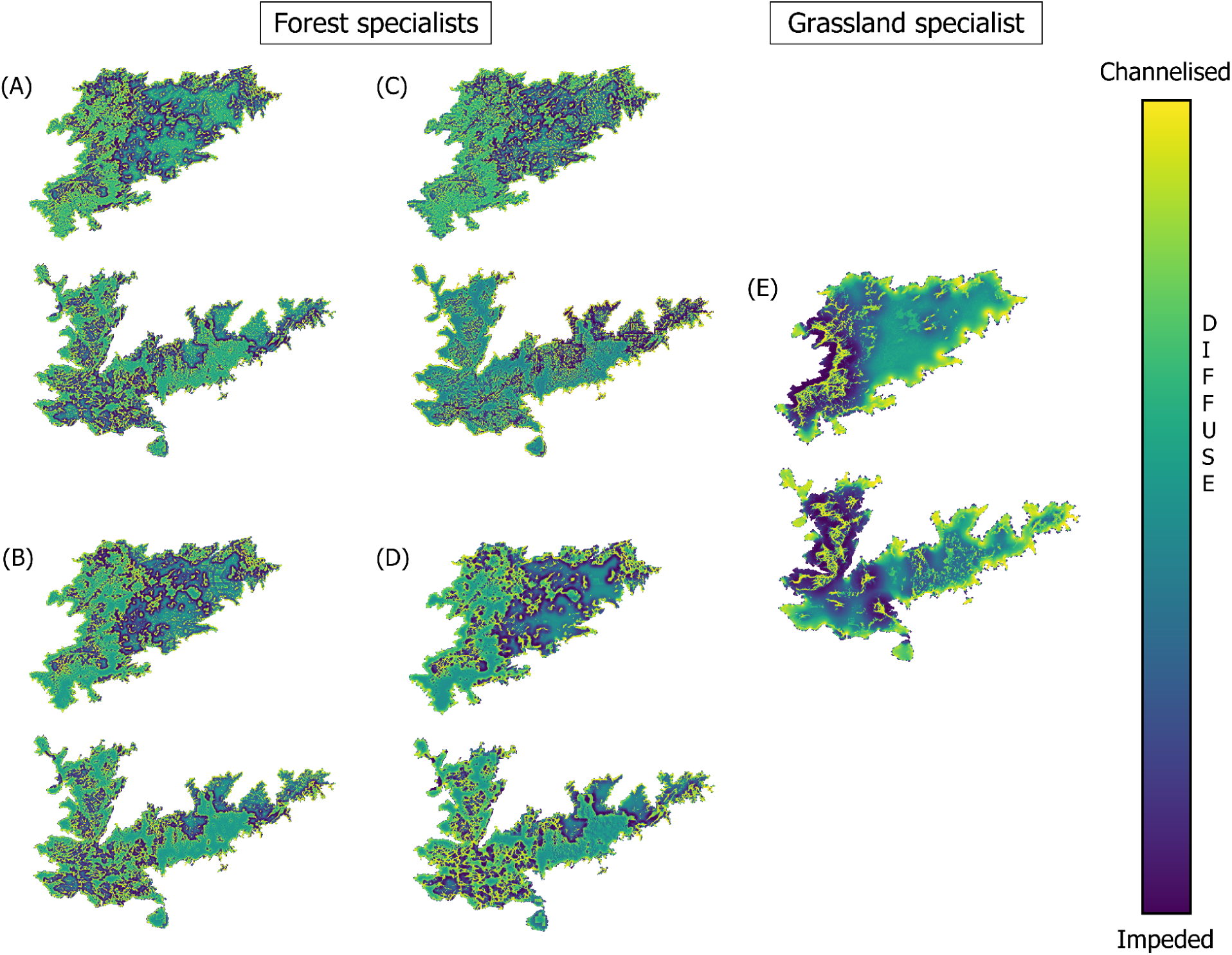
Present normalised current flow (NCF) maps for species A) *Sholicola major* and *Sholicola albiventris* (c = 8, d = 500m), B) *Ficedula nigrorufa* (c = 8, d = 500 m), C) *Montecincla cachinnans* and *Montecincla fairbanki* (c = 8, d = 500m), D) *Eumyias albicaudatus* (c = 8, d = 1000m), E) *Anthus nilghiriensis* (c = 0.25, d = 3000m). *c* determines the non-linearity of the habitat suitability to resistance transformation used to generate the resistance layer, and *d* represents the radius of the moving window in the connectivity model. Colour scale for visualisation is according to percentile bins for NCF values. NCF > 1 represents channelised current flow or potential movement corridors (yellow colours), NCF < 1 represents impeded current flow or potential barriers to movement (dark blue colours), and NCF ≈ 1 represents diffuse current flow (green-ish blue colours), i.e., neither impeded nor channelised. For maps of the other models, refer to Section S2 in the Supplementary Material.

NCF (normalised current flow) is close to 1, across most of the Palani-Anamalai region and the western half of the Nilgiris region for all forest species (Fig. 2), indicating a mostly connected landscape. A small portion of the easternmost end of the Palani-Anamalai hills (slice 5 in Fig. 3), however, seems to be more isolated from the rest of the landscape (Fig. 2). This could be owed to topography and high modification just west of a narrow patch of native forest (locally referred to as the Tiger Shola, located in the Dindigul district), between slice 4 and 5 in Fig. 3. This could result in this patch being a pinch point for movement between two portions of the Palani-Anamalai region (slice 1-4 and slice 5) for forest species. There is also a general declining pattern for cumulative current flow (CCF) going from the west to the east in both regions, with a sharper dip in the Nilgiris, for all forest species.

**Fig. 3:**
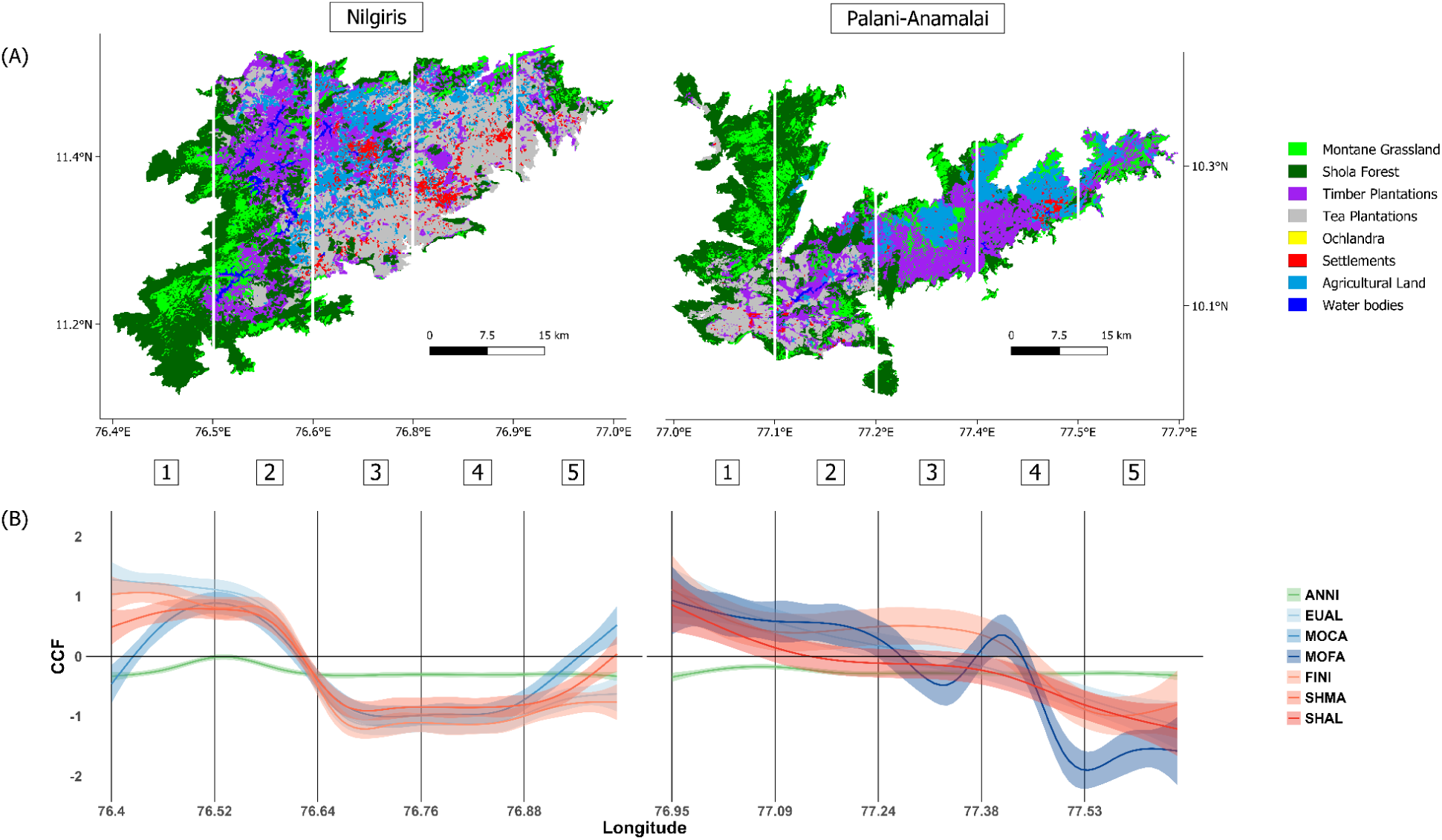
A) Present land use (2017) maps represented in 8 categories for the two Sky Islands. **B) Present cumulative current flow (CCF)** transformed to represent multiples of standard deviations away from the mean for each species over a longitudinal gradient, from west to east on the x-axis for each corresponding Sky Island. Standard deviation stretched values for CCF are plotted with a 95% confidence interval after smoothing using a Generalised Additive Model (GAM). Species key: ANNI - *Anthus nilghiriensis* (grassland specialist), EUAL - *Eumyias albicaudatus* (forest specialist), MOCA - *Montecincla cachinnans* (forest specialist), MOFA - *Montecincla fairbanki* (forest specialist), FINI - *Ficedula nigrorufa* (forest specialist), SHMA - *Sholicola major* (forest specialist), SHAL - *Sholicola albiventris* (forest specialist). For plots of the other models, refer to Section S2 in the Supplementary Material.

Current flow for the grassland species, however, seems to be much more impeded (NCF < 1), with wider barriers (Fig. 2), and lower CCF (Fig. 3) across most of the landscape.

### Change in movement suitability vs. landscape change

We find that current flow seems to follow patterns of landscape change, especially in areas where grassland has been converted to woodlands (Fig. 4A), which represents most of the areas that have been modified (Fig. 4B). There is an overall increase in CCF for the forest species where grassland has been converted to woodland and a decrease for the grassland species within the same category. The only areas where CCF seems to have decreased for forest species is where woodland has been converted into human-use landscapes (Fig. 4A), although this category accounts for a very small percentage of the total area (Fig. 4B). The quantity of grassland gain from other land cover types is negligible, hence it is not feasible to make correlations of grassland gain with change in CCF from these regions. There seems to be no discernible pattern of change in areas where the landscape has not changed significantly during this approximately 20-year period (Fig. 4A). This includes a large portion of the eastern half of the Nilgiris, despite its historic modification, and where woodland and grassland cover have remained stable in the western portions of both landscapes (Fig. 4B, Fig. 5A).

**Fig. 4:**
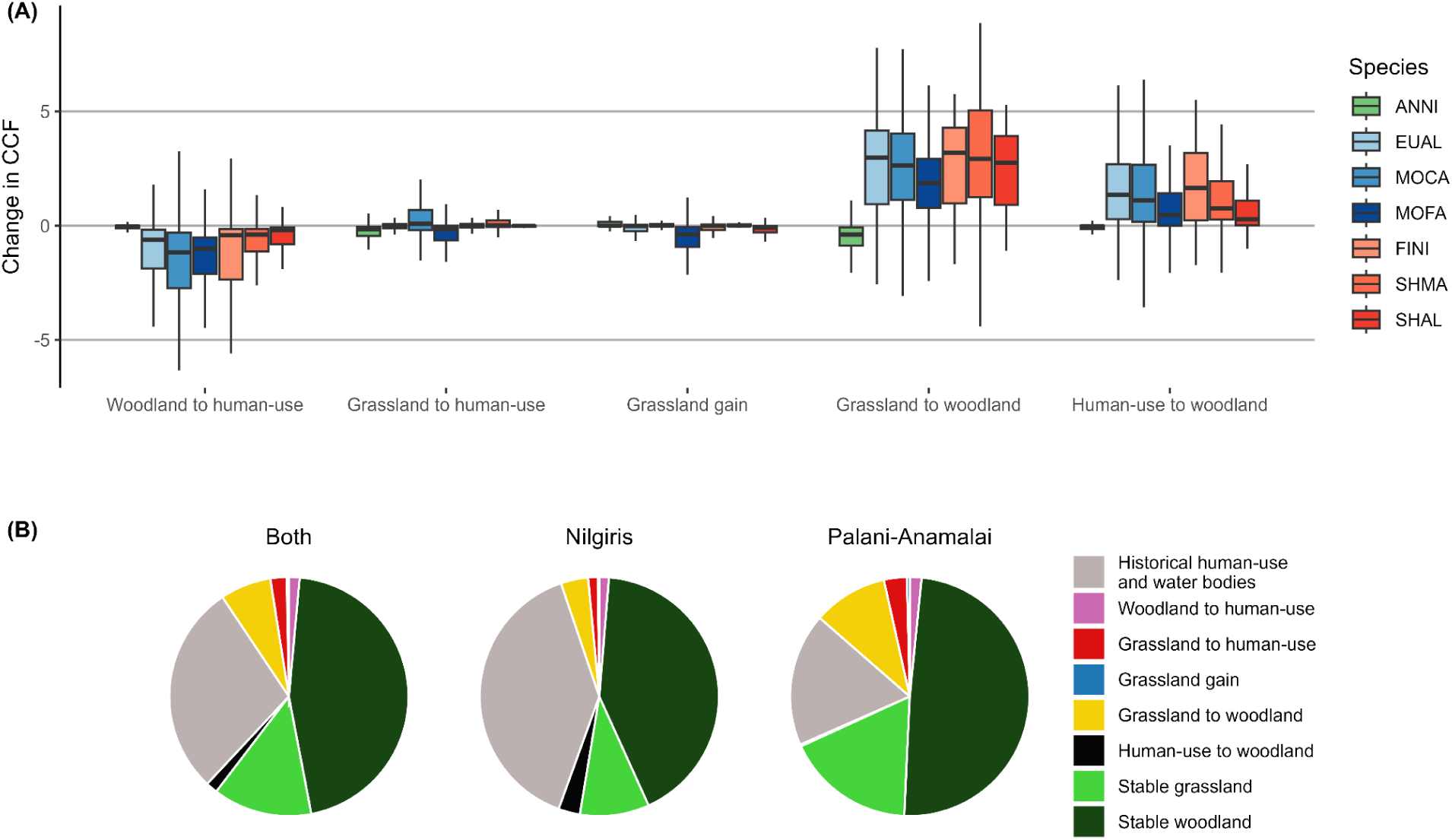
A) Change in cumulative current flow (CCF) within 5 landscape change categories for different species. Values for change in CCF for each species have been transformed to represent multiples of standard deviations away from zero. Change in CCF > 0 represents an increase, change in CCF < 0 represents a decrease, and change in CCF ≈ 0 represents no change in current flow (movement suitability). Species key: ANNI - *Anthus nilghiriensis* (grassland specialist), EUAL - *Eumyias albicaudatus* (forest specialist), MOCA - *Montecincla cachinnans* (forest specialist), MOFA - *Montecincla fairbanki* (forest specialist), FINI - *Ficedula nigrorufa* (forest specialist), SHMA - *Sholicola major* (forest specialist), SHAL - *Sholicola albiventris* (forest specialist). Refer to Section S3 in the Supplementary Material for the complete figure, including all landscape change categories and outlier values, and plots with all models for each species. B) Fraction of total area covered by each land use change category across both sky islands and on each individual sky island.

**Fig. 5:**
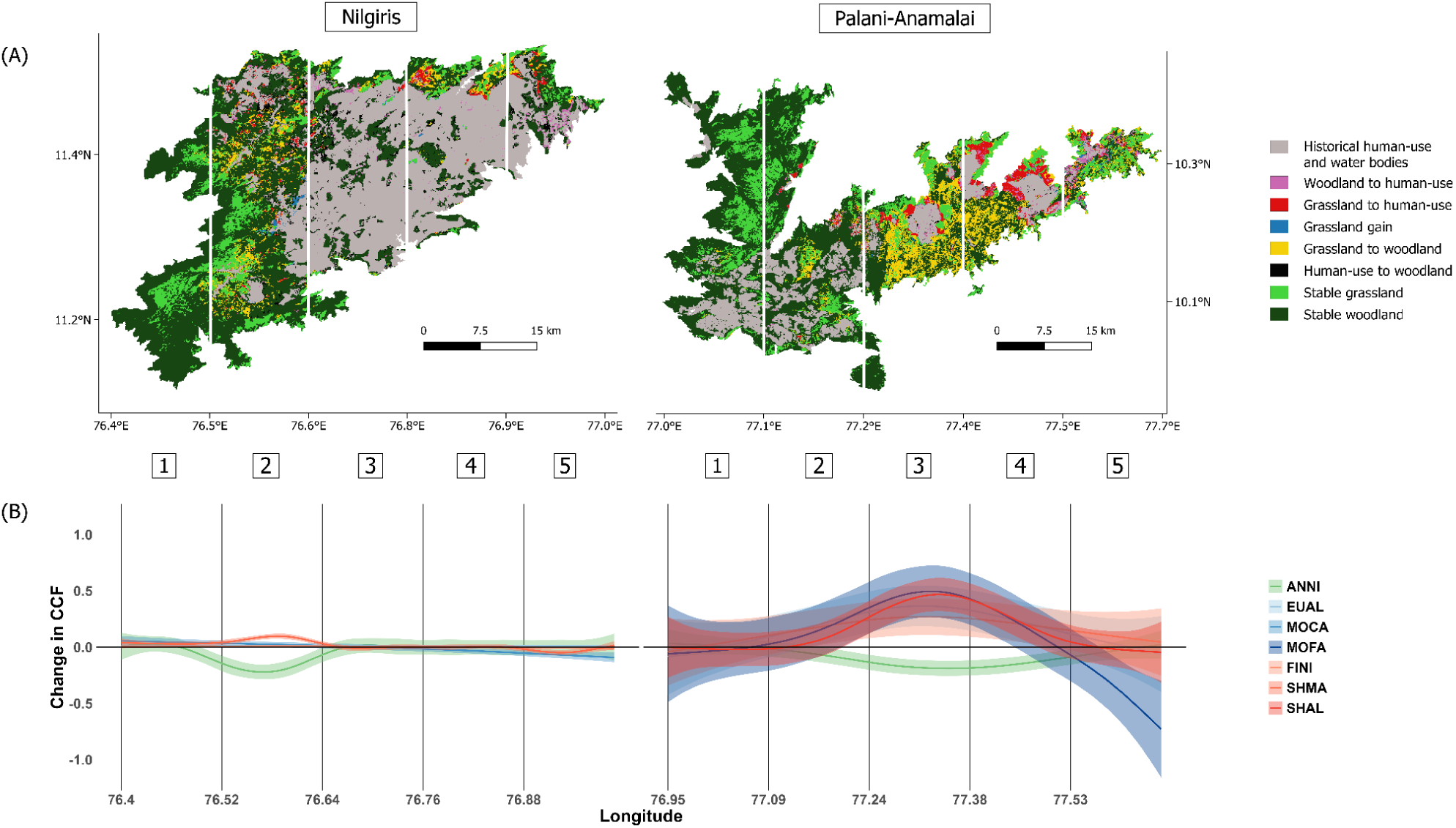
A) Landscape change (from 1995 to 2017) maps represented in 8 categories for the two Sky Islands. **B) Change in cumulative current flow (CCF)** transformed to represent multiples of standard deviations away from zero for each species over a longitudinal gradient, from west to east on the x-axis, for each corresponding Sky Island. Change in CCF > 0 represents an increase, change in CCF < 0 represents a decrease, and change in CCF ≈ 0 represents no change in current flow (movement suitability). Standard deviation stretched values for change in CCF are plotted with a 95% confidence interval after smoothing using a Generalised Additive Model (GAM). Species key: ANNI - *Anthus nilghiriensis* (grassland specialist), EUAL - *Eumyias albicaudatus* (forest specialist), MOCA - *Montecincla cachinnans* (forest specialist), MOFA - *Montecincla fairbanki* (forest specialist), FINI - *Ficedula nigrorufa* (forest specialist), SHMA - *Sholicola major* (forest specialist), SHAL - *Sholicola albiventris* (forest specialist). For plots and maps of the other models, refer to Section S4 in the Supplementary Material.

### Species-specific changes in spatial patterns of movement

We do not recover a differential change in CCF between different forest species (Fig. 5B). However, there seems to be a large difference in the change in CCF between forest and grassland species. Change in CCF appears to follow a flipped pattern for the grassland species, having reduced in the same regions where it has increased for the forest species (Fig. 5B).

However, this pattern is almost negligible in the Nilgiris region as compared to the Palani-Anamalai region (see Discussion for details), where the increase in CCF for forest species is also a lot more than the decrease for grassland species.

There does not seem to be a large difference in the actual patterns of change in current flow and present current flow between models using different dispersal distance thresholds and resistance layers. However, there might be a difference in the intensity of those patterns across different models, especially with different *c* values, increasing in intensity with more lenient (non-linear) habitat suitability to movement resistance transformations, i.e., higher *c* values (refer to Figures S5 to S22 in the Supplementary Material).

## Discussion

Our study provides insight into patterns of habitat connectivity for six forest and one grassland specialist species. It covers two of the largest Sky Islands in the Western Ghats of India, comprising most of the global distribution of five species (*Eumyias albicaudatus*, *Montecincla fairbanki*, *Ficedula nigrorufa*, *Sholicola major* and *Sholicola albiventris*), and the entire global distribution of two species (*Anthus nilghiriensis* and *Montecincla cachinnans*). Specifically, it provides valuable insights into how connectivity may have changed for species with different habitat requirements over approximately two decades. It identifies landscape change, especially the invasion of grasslands by woodlands, as a major driver of these changes.

We find a general decreasing trend in movement suitability going from the east to the west, which might impact connectivity between populations in the east and the west. Genomic differentiation between populations has been attributed mostly to barriers for gene flow by climate or landscape features (Lee & Mitchell-Olds, 2011; Yang et al., 2013), often along latitudinal (Zakharov & Hellmann, 2008) or longitudinal gradients (Geffen et al., 2004). The spatial patterns of movement suitability for our study species seem to corroborate a pattern of separation of bioclimatic zones along an east-west axis in the region (Pascal, 1982; Sekar & Karanth, 2013). Speciation, across this longitudinal climatic gradient, of a species of frog, *Nasikabatrachus bhupathi*, has been ascribed to monsoonal patterns (Janani et al., 01 2017). However, in our case, the gradient in movement suitability might also be attributed to patterns of land use, with there being an east-west gradient in anthropogenic modification on both Shola Sky Islands (Arasumani et al., 2019; Joshi et al., 2018). Their western halves are largely comprised of designated protected areas (Arasumani et al., 2019) that preserve the natural stable biphasic structure of the landscape, thus facilitating movement for both kinds of habitat specialists. Much of the modification is concentrated in the eastern halves of both Shola Sky Islands. Our study highlights the importance of protecting natural landscapes to conserve habitat connectivity for multiple species. Protected areas can act as important stepping stones and corridors to preserve habitat connectivity between populations in disturbed landscapes (Luja et al., 2017; Schultz, 1998).

Species responses to landscape change, however, vary depending on the type and extent of change (Rivera-Ortíz et al., 2015) as well as species ecology. Factors such as mobility (Rivera-Ortíz et al., 2015), type and degree of habitat specialisation (Coppedge et al., 2001; Jobin et al., 2025), and niche breadth (Peyras et al., 2013; Wehner et al., 2021) could influence species responses to landscape change. In this region, we know that while both forest and grassland specialists in this region show an avoidance at varying levels to purely human-use landscapes, such as tea plantations, settlements and other agricultural land, their responses to wooded habitats are contrasting (Jobin et al., 2025). This seems to be reflected in the differences in patterns of movement between the two Sky Islands. Most of the eastern half of the Nilgiris has been transformed for tea production, with very little woodland cover for over 150 years (Joshi et al., 2018; Ramesh et al., 2025). However, a large portion of the eastern half of the Palani-Anamalais is woodland, either naturally or through invasion. Trends in movement suitability along an east-west axis seem to track these patterns. For the forest specialists, there is a sharp decrease in the Nilgiris and a more gradual decrease in the Palani-Anamalai region from the west to the east. However, the grassland species, with its stricter habitat requirements, show similar patterns in both landscapes along an east-west axis.

Our results indicate that these landscape modifications will have strong negative consequences for the grassland species in this region; woodland habitat connectivity seems to have improved overall at the cost of grassland habitat connectivity over approximately two decades. This is in line with global trends in the loss and increased fragmentation of open habitats over the last century (Bardgett et al., 2021; Bonanomi et al., 2019; Parr et al., 2014), and a subsequent decline in open habitat species populations (Knopf, 1994; Prangel et al., 2023). Contemporary genetic differentiation between populations in other highly specialised species reflects recent landscape modifications as opposed to long-term historical changes (da Silva Carvalho et al., 2015; Robin, Gupta, et al., 2015; Vandergast et al., 2007; Zellmer & Knowles, 2009). Hence, this reduction in grassland habitat, especially in the middle regions of the Palani-Anamalai hills (Arasumani et al., 2019), might impact gene flow and subsequently genetic differentiation between grassland species populations in the east and the west in the near future. On the other hand, forest specialists may have benefited from this kind of landscape change. Non-native woodlands seem to have filled in gaps between native forests, resulting in larger, more contiguous patches of suitable woodland habitat and thus improved woodland habitat connectivity.

### Conclusions and future directions

Our study highlights the effects of both the type of anthropogenic modification and species ecology on habitat connectivity. While the general trend is to expect anthropogenic modification of natural landscape structures to impact species negatively, our study indicates that species occupying habitats in close proximity to each other can vary widely in their response to landscape change, depending on their specialisation.

Despite the clear direction of our results, there are several challenges in working with wild birds in remote mountain-tops and subsequent limitations with the available information on species ecology. Data on dispersal behaviour is especially not feasible to collect given the recent changes in the legal provisions in obtaining permits for capture and release of birds (Shanker et al., 2023) as well as historic challenges in conducting fieldwork in these regions (Madhusudan et al., 2006). A better understanding of species ecology may, however, help to predict more precisely species response to future landscape change scenarios. Future studies exploring genetic connectivity between populations across this landscape could also give further insight into the effects of the loss and gain in habitat connectivity due to landscape change on these populations.

## Acknowledgements

This work was supported by the National Geographic Society Level-II grant (NGS-93271R-22), and Rohini Nilekani Philanthropies (G-202410-00940). We thank Akshay Herur for generating the climate zone maps. We thank Devcharan Jathanna for his insights that helped construct this study, and Jan Engler and Trevor Price for discussions at various stages of the project. We also thank Ashwin Warudkar, Naman Goyal, Viral Joshi and other lab members at IISER Tirupati for detailed discussions throughout the duration of the project.

## CRediT authorship contribution statement

**Arunima Jain:** Conceptualisation, Data curation, Formal analysis, Investigation, Methodology, Software, Visualisation, Writing - original draft, Writing - review & editing. **Chiti Arvind:** Conceptualisation, Data curation, Formal analysis, Investigation, Methodology, Writing - review & editing. **Jobin Varughese:** Conceptualisation, Data curation, Formal analysis, Investigation, Methodology, Writing - review & editing. **Abhimanyu Lele:** Investigation, Methodology, Writing - review & editing. **V.V. Robin:** Conceptualisation, Funding acquisition, Investigation, Methodology, Project administration, Resources, Supervision, Writing - original draft, Writing - review & editing.

## Notes

### Competing Interest Statement

The authors have declared no competing interest.

