## Supplementary Material for "Changes in habitat connectivity for range-restricted birds reflect patterns of woodland invasion"

**S1. Species Distribution Models**

**Table S1:** Predictor variables used for species distribution models

| Type | Variable name | Description | Source (present) | Source (past) | Original resolution |
|---|---|---|---|---|---|
| Landscape | dist_shola (only used for forest species) | Distance from nearest Shola forest edge in metres | Derived from 2017 land cover map (Arasumani et al. 2019) using raster proximity function in QGIS 3.28.11 | Derived from 1995 land cover map (Arasumani et al. 2019) using raster proximity function in QGIS 3.28.11 | 30 m x 30 m |
| Landscape | dist_woodland (negatively correlated with Woodland_cover for Palani-Anamalai region and positively with dist_shola for Nilgiris region) | Distance from nearest woodland edge in metres | Derived from 2017 land cover map (Arasumani et al. 2019) using raster proximity function in QGIS 3.28.11 | Derived from 1995 land cover map (Arasumani et al. 2019) using raster proximity function in QGIS 3.28.11 | 30 m x 30 m |
| Landscape | dist_settlements (only used for forest species) | Distance from nearest settlement in metres | Derived from 2017 land cover map (Arasumani et al. 2019) using raster proximity function in QGIS 3.28.11 | Derived from 1995 land cover map (Arasumani et al. 2019) using raster proximity function in QGIS 3.28.11 | 30 m x 30 m |
| Landscape | dist_grassland (only used for grassland species) | Distance from nearest grassland edge in metres | Derived from 2017 land cover map (Arasumani et al. 2019) using raster proximity function in QGIS 3.28.11 | Derived from 1995 land cover map (Arasumani et al. 2019) using raster proximity function in QGIS 3.28.11 | 30 m x 30 m |

| Type | Variable name | Description | Source (present) | Source (past) | Original resolution |
| --- | --- | --- | --- | --- | --- |
| Landscape | Shola_Forest_cover (only used for forest species) | Percentage cover of Shola Forest | Derived from 2017 polygonised land cover map (Arasumani et al. 2019) using *terra 1.7-71* package in R | Derived from 1995 polygonised land cover map (Arasumani et al. 2019) using *terra 1.7-71* package in R | 30 m x 30 m |
| Landscape | Woodland_cover (only used for forest species) | Percentage cover of Woodland (Shola forest + Timber plantations) | Derived from 2017 polygonised land cover map (Arasumani et al. 2019) using *terra 1.7-71* package in R | Derived from 1995 polygonised land cover map (Arasumani et al. 2019) using *terra 1.7-71* package in R | 30 m x 30 m |
| Landscape | Timber_Plantations_cover (only used for forest species) | Percentage cover of Timber plantations | Derived from 2017 polygonised land cover map (Arasumani et al. 2019) using *terra 1.7-71* package in R | Derived from 1995 polygonised land cover map (Arasumani et al. 2019) using *terra 1.7-71* package in R | 30 m x 30 m |
| Landscape | Grassland_cover (only used for grassland species) | Percentage cover of Montane Grassland | Derived from 2017 polygonised land cover map (Arasumani et al. 2019) using *terra 1.7-71* package in R | Derived from 1995 polygonised land cover map (Arasumani et al. 2019) using *terra 1.7-71* package in R | 30 m x 30 m |
| Topography | elevation (only used for grassland | Elevation in metres ASL | NASA Shuttle Radar Topography | Same as for present | 30 m x 30 m |

| Type | Variable name | Description | Source (present) | Source (past) | Original resolution |
|---|---|---|---|---|---|
| | species) | | Mission DEM Global 1 Arc Second V003 | | |
| Topography | slope (correlated to TRI) | Slope in degrees | Derived from DEM using *terra 1.7-71* package in R | Same as for present | 30 m x 30 m |
| Topography | aspect | Aspect in degrees | Derived from DEM using *terra 1.7-71* package in R | Same as for present | 30 m x 30 m |
| Topography | roughness (correlated to TRI) | Roughness in metres | Derived from DEM using *terra 1.7-71* package in R | Same as for present | 30 m x 30 m |
| Topography | TRI | Terrain Ruggedness Index in metres | Derived from DEM using *terra 1.7-71* package in R | Same as for present | 30 m x 30 m |
| Topography | TPI | Topographic Position Index in metres | Derived from DEM using *terra 1.7-71* package in R | Same as for present | 30 m x 30 m |
| Topography | TWI | Topographic Wetness Index | Derived from DEM using SAGA Next Gen in QGIS 3.28.11 | Same as for present | 30 m x 30 m |
| Vegetation | canopyheight | Canopy height in metres | Year 2020 dataset from Potapov et al. (2021) | Year 2000 dataset from Potapov et al. (2021) | 30 m x 30 m |
| Climate | clim_zone | Categorical zones derived from remotely sensed Precipitation and Mean daily air | (Pascal 1982; Karger et al. 2017; Karger et al. 2021) | Same as for present | 1000 m x 1000 m |

| Type | Variable name | Description | Source (present) | Source (past) | Original resolution |
|---|---|---|---|---|---|
| | | temperature (TAS) data based on Pascal (1982)'s bioclimatic zones for the Western Ghats | | | |

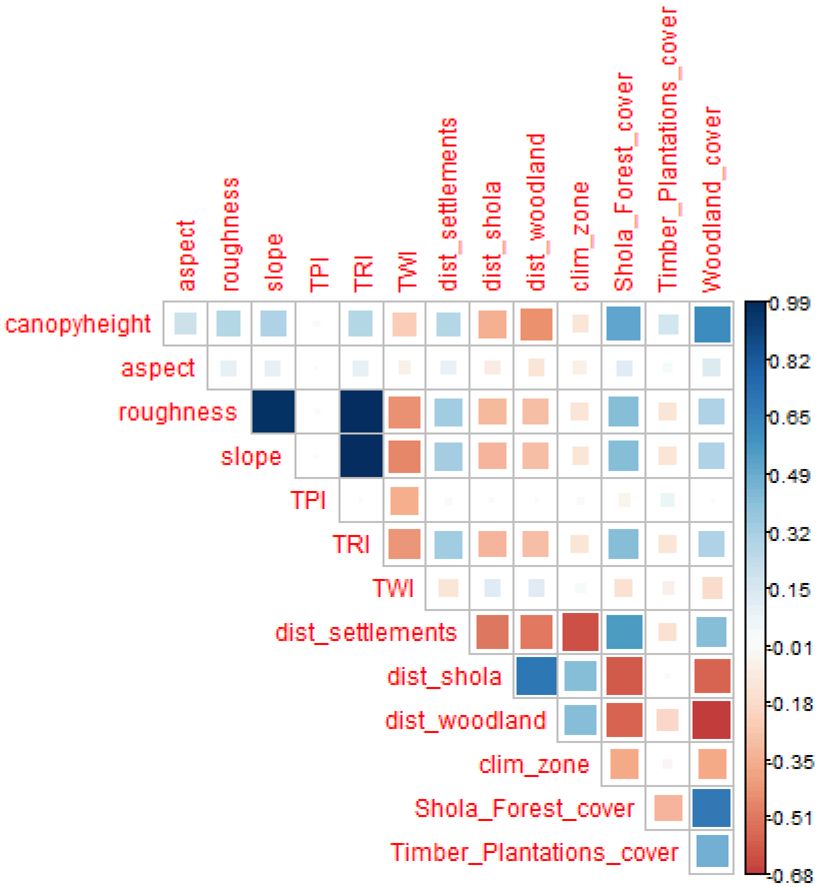


Fig. S1: Pearson's correlation index matrix for predictor variables for the Nilgiris

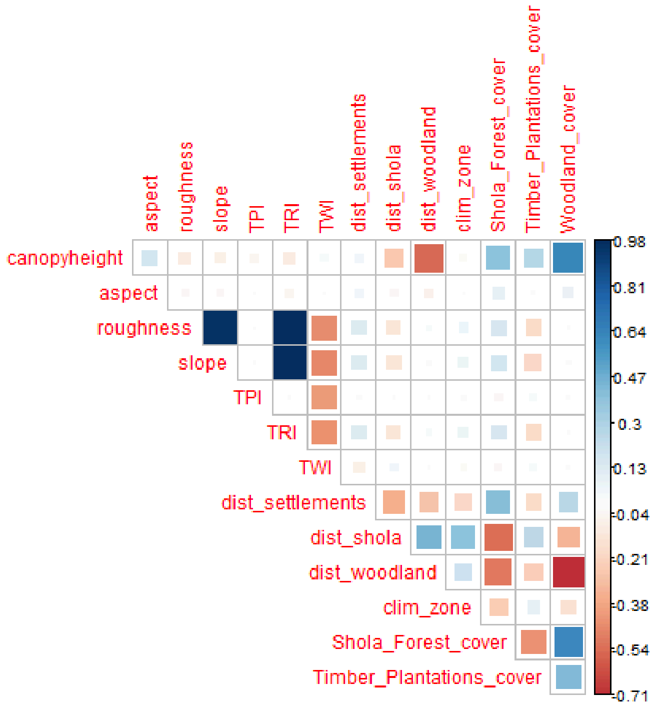


Fig. S2: Pearson's correlation index matrix for predictor variables for the Palani-Anamalai region

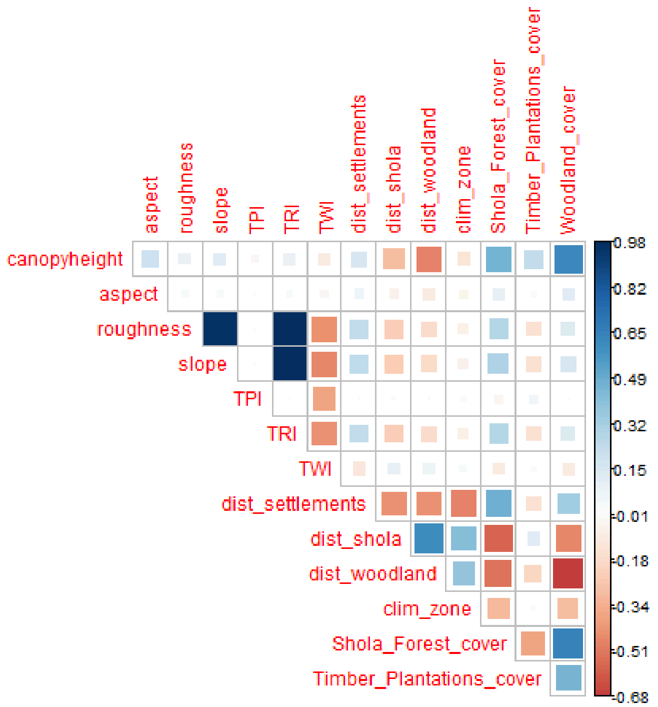


Fig. S3: Pearson's correlation index matrix for predictor variables for forest species for both regions together

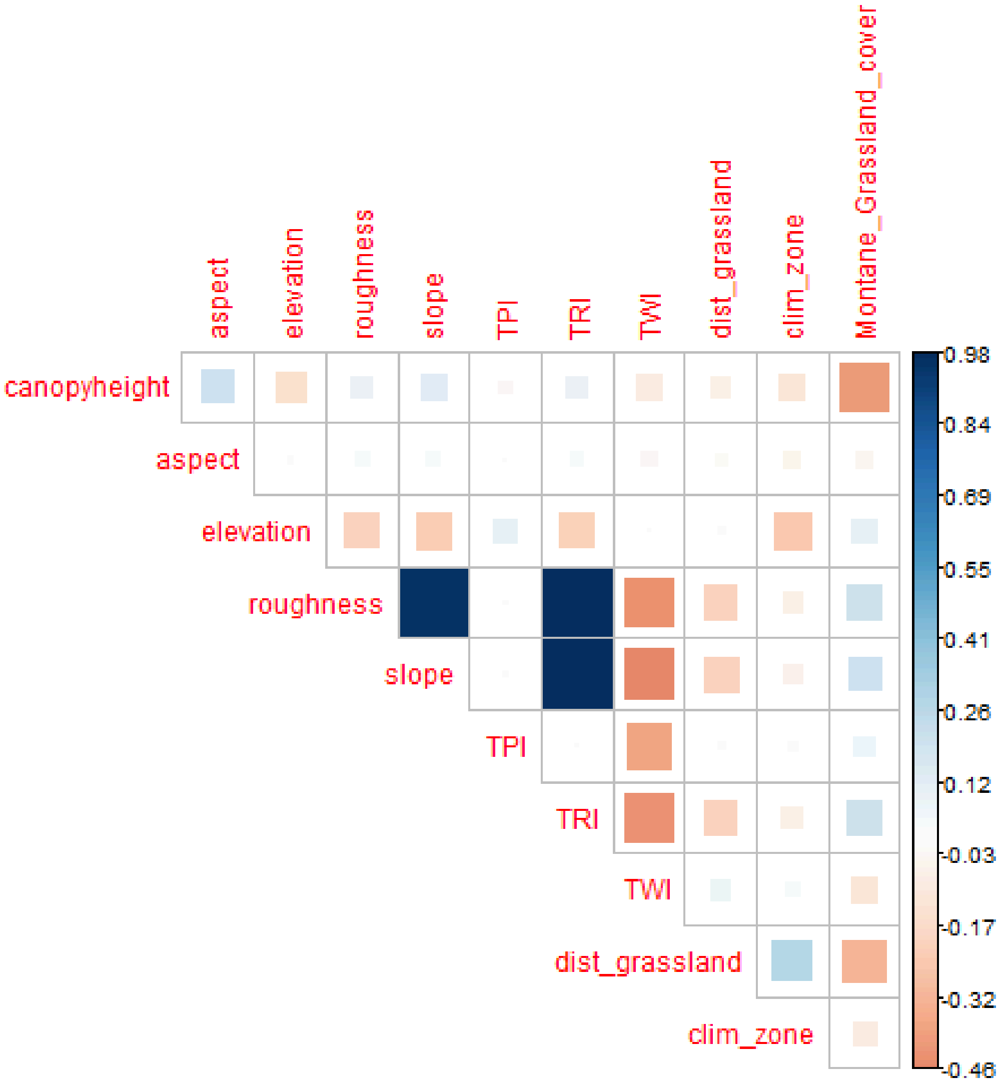


Fig. S4: Pearson’s correlation index matrix for predictor variables for grassland species for both regions together

**Table S2:** Data used to make species distribution maps for each species along with model evaluation scores and hyperparameters (mtry - number of predictors sampled at each node, nodesize - minimum size of terminal nodes (Liaw and Wiener 2002)) Hyperparameters were tuned using the *tuneRanger* package in R (Probst et al. 2018). Species key: ANNI - *Anthus nilghiriensis* (grassland specialist), EUAL - *Eumyias albicaudatus* (forest specialist), MOCA - *Montecincla cachinnans* (forest specialist), MOFA - *Montecincla fairbanki* (forest specialist), FINI - *Ficedula nigrorufa* (forest specialist), SHMA - *Sholicola major* (forest specialist), SHAL - *Sholicola albiventris* (forest specialist).

| | **SHAL** | **SHMA** | **MOFA** | **MOCA** | **ANNI** | **EUAL** | **FINI** |
|---|---|---|---|---|---|---|---|
| **Data type** | Presence /Absence | Presence /Absence | Presence /Absence | Presence /Absence | Presence/ Pseudo-absence | Presence/ Absence | Presence/ Absence |
| **Region** | Palani-Anamalais | Nilgiris | Palani-Anamalais | Nilgiris | Both | Both | Both |
| **Presences (train)** | 175 | 112 | 215 | 122 | 593 | 313 | 371 |
| **Presences (test)** | 41 | 20 | 51 | 32 | 151 | 77 | 100 |
| **Absences (train)** | 121 | 126 | 89 | 126 | 4003 | 223 | 183 |
| **Absences (test)** | 32 | 39 | 24 | 30 | 997 | 57 | 38 |
| **AUC (ROC)** | 0.870427 | 0.857692 | 0.707925 | 0.764062 | 0.98354 | 0.776942 | 0.801579 |
| **AUC (PRC)** | 0.79663 | 0.743881 | 0.850463 | 0.782579 | 0.901117 | 0.83025 | 0.86171 |
| **mtry** | 3 | 3 | 4 | 1 | 3 | 8 | 6 |
| **nodesize** | 27 | 10 | 61 | 3 | 2 | 106 | 58 |

**Table S3:** Variable importance measures for each species based on mean decrease in accuracy (not standardised across models). Species key: ANNI - *Anthus nilghiriensis* (grassland specialist), EUAL - *Eumyias albicaudatus* (forest specialist), MOCA - *Montecincla cachinnans* (forest specialist), MOFA - *Montecincla fairbanki* (forest specialist), FINI - *Ficedula nigrorufa* (forest specialist), SHMA - *Sholicola major* (forest specialist), SHAL - *Sholicola albiventris* (forest specialist).

| | SHAL | SHMA | MOFA | MOCA | ANNI | EUAL | FINI |
|---|---|---|---|---|---|---|---|
| **canopyheight** | 10.17998 | 18.73652 | 4.862911 | 14.27173 | 48.33227 | 4.994026 | 21.69257 |
| **aspect** | 3.271299 | -3.8814 | -5.57708 | -0.52427 | 11.91337 | 4.27143 | 17.07446 |
| **TPI** | 4.416082 | 2.87834 | 2.718836 | 0.593759 | 9.037364 | -0.86096 | 3.613103 |
| **TRI** | 0.746168 | 2.612128 | 0.355272 | 11.13325 | 35.59626 | 0.105439 | 6.844347 |
| **TWI** | -0.40501 | -0.79114 | 5.014464 | 7.003905 | 9.812246 | -2.18996 | 1.384094 |
| **dist_settlements** | 9.717539 | 16.67125 | 5.126851 | 17.52399 | NA | 4.875014 | 4.530247 |
| **dist_shola** | 14.55021 | 15.50235 | 16.68773 | 18.92544 | NA | 12.72184 | 8.306876 |
| **clim_zone** | 36.29363 | 19.10435 | 26.49466 | 16.1158 | 67.50337 | 35.90642 | 54.60974 |
| **Shola_Forest_cover** | 11.85107 | 8.716857 | 10.18061 | 17.90746 | NA | 2.619006 | 5.548079 |
| **Timber_Plantations_cover** | 8.875369 | 11.74991 | 6.852745 | 15.56083 | NA | 2.850253 | 8.062539 |
| **Woodland_cover** | 31.18257 | 33.31216 | 9.475409 | 23.99214 | NA | 36.95521 | 52.41349 |
| **dist_woodland** | NA | NA | NA | NA | NA | 17.28714 | 21.05343 |
| **elevation** | NA | NA | NA | NA | 38.95881 | NA | NA |
| **dist_grassland** | NA | NA | NA | NA | 28.90343 | NA | NA |
| **Montane_Grassland_cover** | NA | NA | NA | NA | 21.12046 | NA | NA |

### S2. Spatial patterns of present connectivity

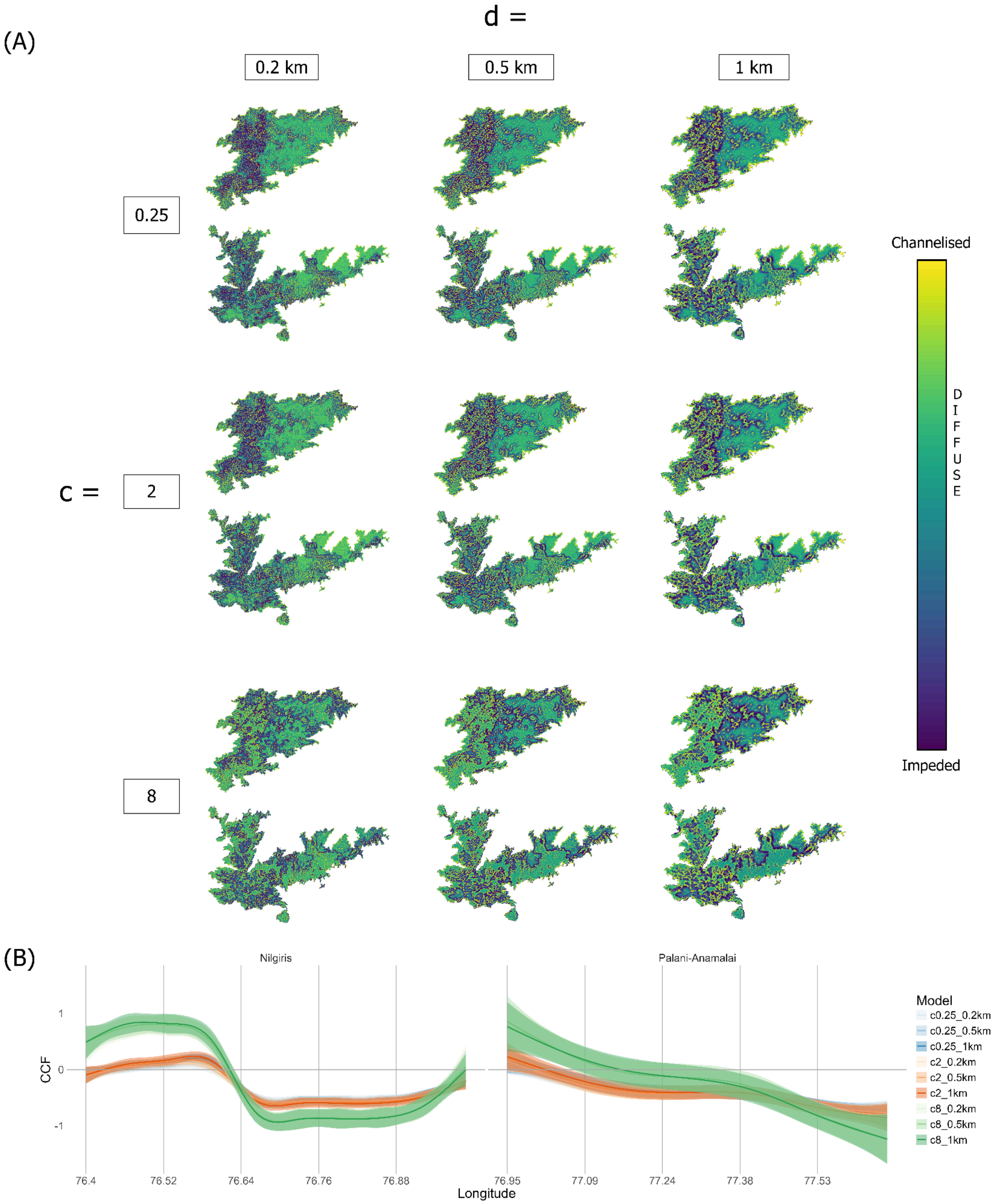


Fig. S5: Species - *Sholicola major* (in the Nilgiris) and *Sholicola albiventris* (in the Palani-Anamalais). A) Present-day normalised current flow (NCF). Colour scale for visualisation is according to percentile bins for NCF values. NCF > 1 represents channelised current flow or potential movement corridors (yellow colours), NCF < 1 represents impeded current flow or potential barriers to movement (dark blue colours), and NCF ≈ 1 represents diffuse current flow (green-ish blue colours), i.e., neither impeded nor channelised. B) Present-day cumulative current flow (CCF)

transformed to represent multiples of standard deviations away from the mean for each model (*c* value x dispersal distance), over a longitudinal gradient going from west to east on the x-axis for each corresponding sky island. Standard deviation stretched values for CCF are plotted with a 95% confidence interval after smoothing using a Generalised Additive Model (GAM). *c* determines the non-linearity of the habitat suitability to resistance transformation used to generate the resistance layer, and *d* represents the radius of the moving window in the connectivity model.

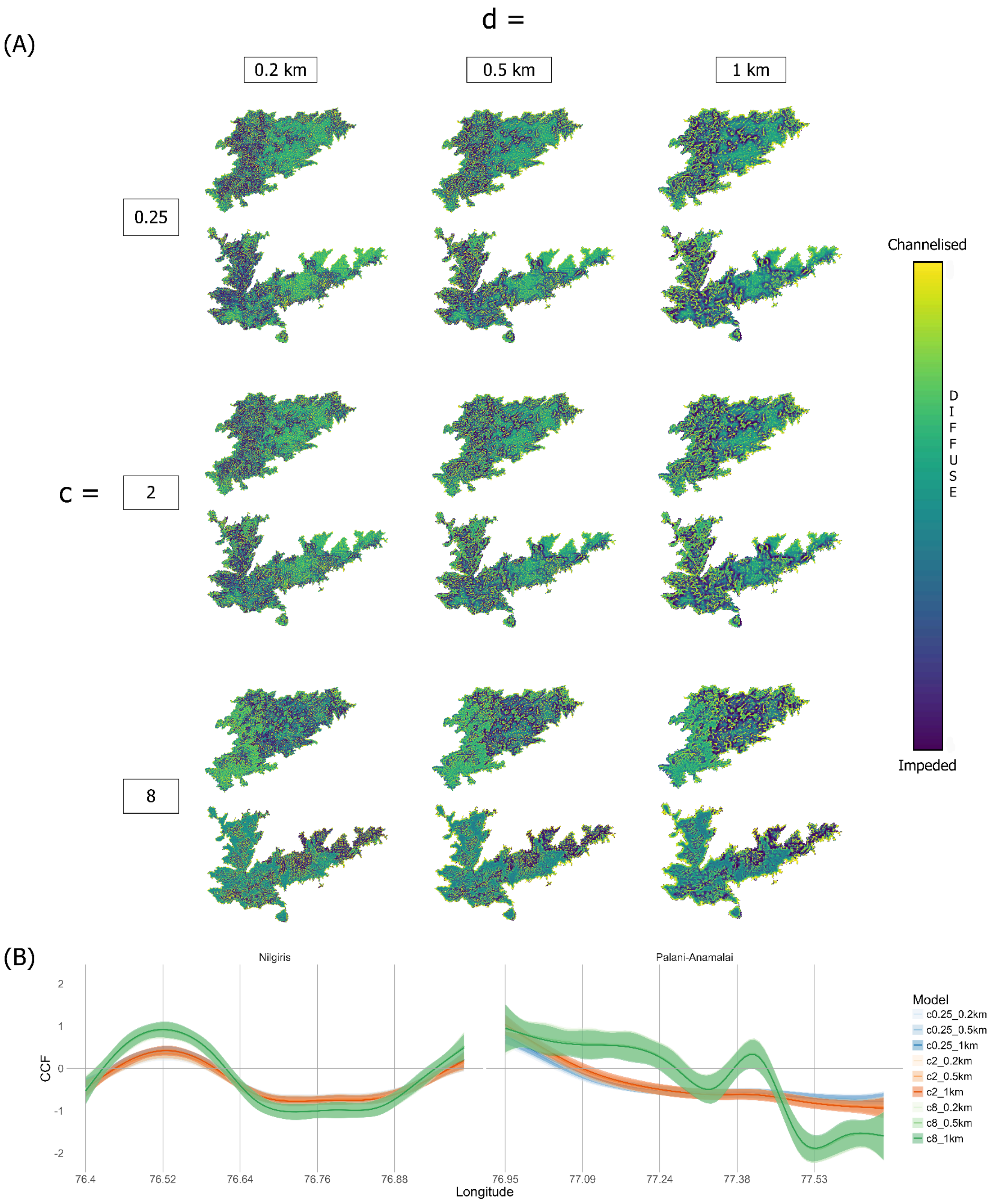


Fig. S6: Species - *Montecincla cachinnans* (in the Nilgiris) and *Montecincla fairbanki* (in the Palani-Anamalais). A) Present-day normalised current flow (NCF). Colour scale for visualisation is according to percentile bins for NCF values. NCF > 1 represents channelised current flow or potential movement corridors (yellow colours), NCF < 1 represents impeded current flow or potential barriers to movement (dark blue colours), and NCF ≈ 1 represents diffuse current flow (green-ish blue colours), i.e., neither impeded nor channelised. B) Present-day cumulative current flow (CCF) transformed to represent multiples of standard deviations away from the mean for each model (*c* value x dispersal distance), over a longitudinal gradient going from west to east on the x-axis for each corresponding sky island. Standard deviation stretched values for CCF are plotted with a 95% confidence interval after smoothing using a Generalised Additive Model (GAM). *c* determines the non-linearity of the habitat suitability to resistance transformation used to generate the resistance layer, and *d* represents the radius of the moving window in the connectivity model.

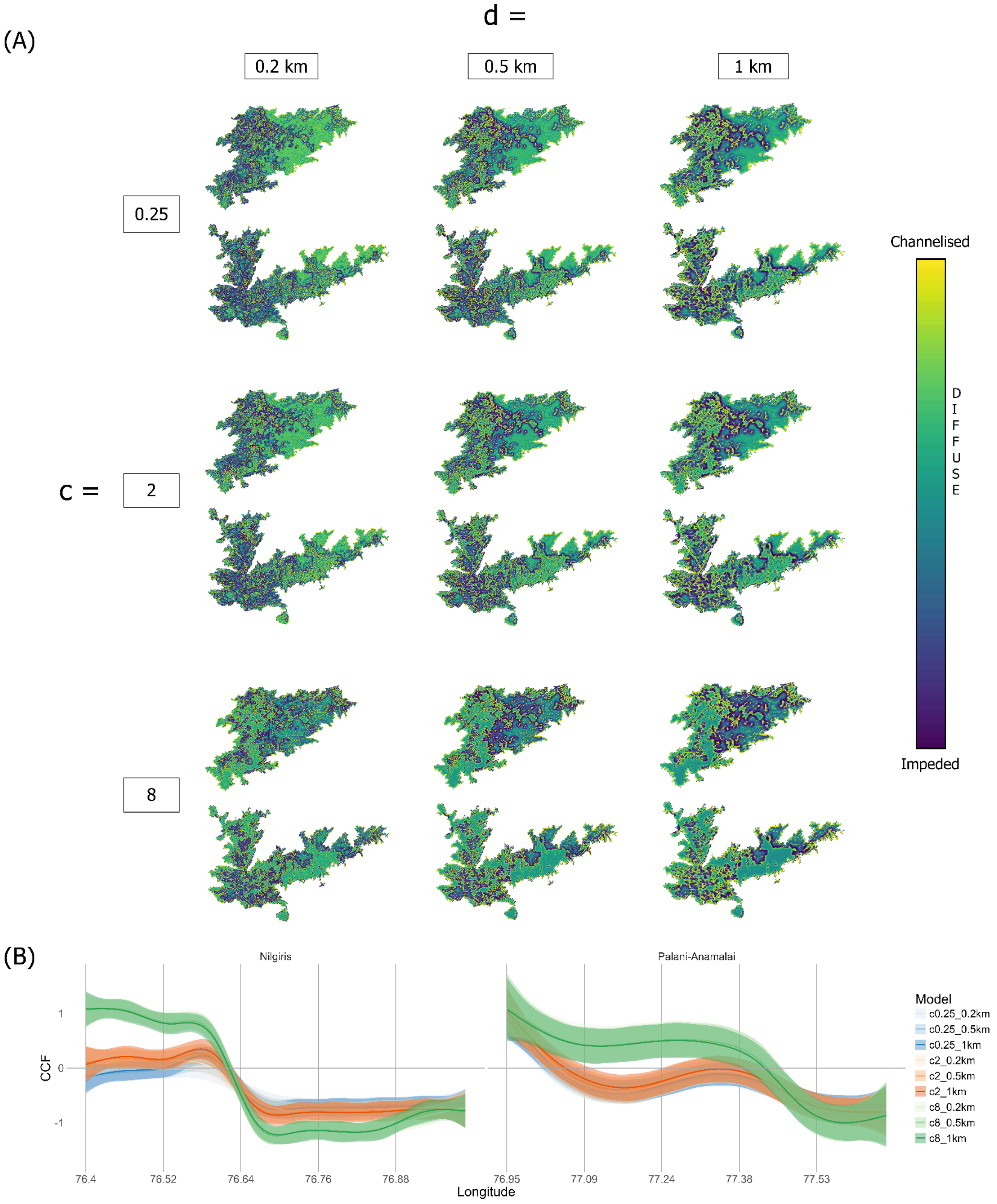


Fig. S7: Species - *Ficedula nigrorufa*. A) Present-day normalised current flow (NCF). Colour scale for visualisation is according to percentile bins for NCF values. NCF > 1 represents channelised current flow or potential movement corridors (yellow colours), NCF < 1 represents impeded current flow or potential barriers to movement (dark blue colours), and NCF ≈ 1 represents diffuse current flow (green-ish blue colours), i.e., neither impeded nor channelised. B) Present-day cumulative current flow (CCF) transformed to represent multiples of standard deviations away from the mean for each model (*c* value x dispersal distance), over a longitudinal gradient going from west to east on the x-axis for each corresponding sky island. Standard deviation stretched values for CCF are plotted with a 95% confidence interval after smoothing using a Generalised Additive Model (GAM). *c* determines the non-linearity of the habitat suitability to resistance transformation used to generate the resistance layer, and *d* represents the radius of the moving window in the connectivity model.

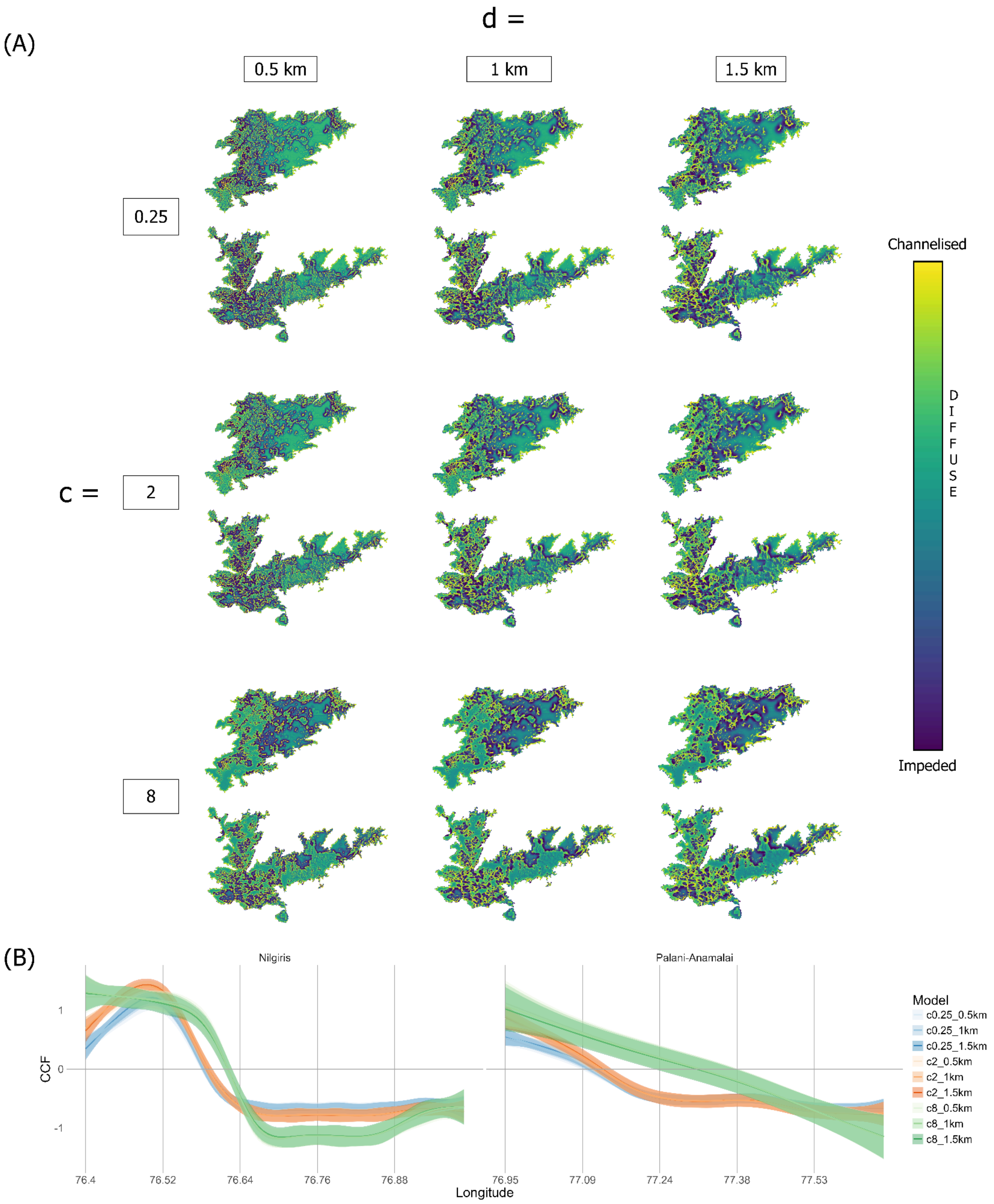


Fig. S8: Species - *Eumyias albicaudatus*. A) Present-day normalised current flow (NCF). Colour scale for visualisation is according to percentile bins for NCF values. NCF > 1 represents channelised current flow or potential movement corridors (yellow colours), NCF < 1 represents impeded current flow or potential barriers to movement (dark blue colours), and NCF ≈ 1 represents diffuse current flow (green-ish blue colours), i.e., neither impeded nor channelised. B) Present-day cumulative current flow (CCF) transformed to represent multiples of standard deviations away from the mean for each model (*c* value x dispersal distance), over a longitudinal gradient going from west to east on the x-axis for each corresponding sky island. Standard deviation stretched values for CCF are plotted with a 95% confidence interval after smoothing using a Generalised Additive Model (GAM). *c* determines the non-linearity of the habitat suitability to resistance transformation used to generate the resistance layer, and *d* represents the radius of the moving window in the connectivity model.

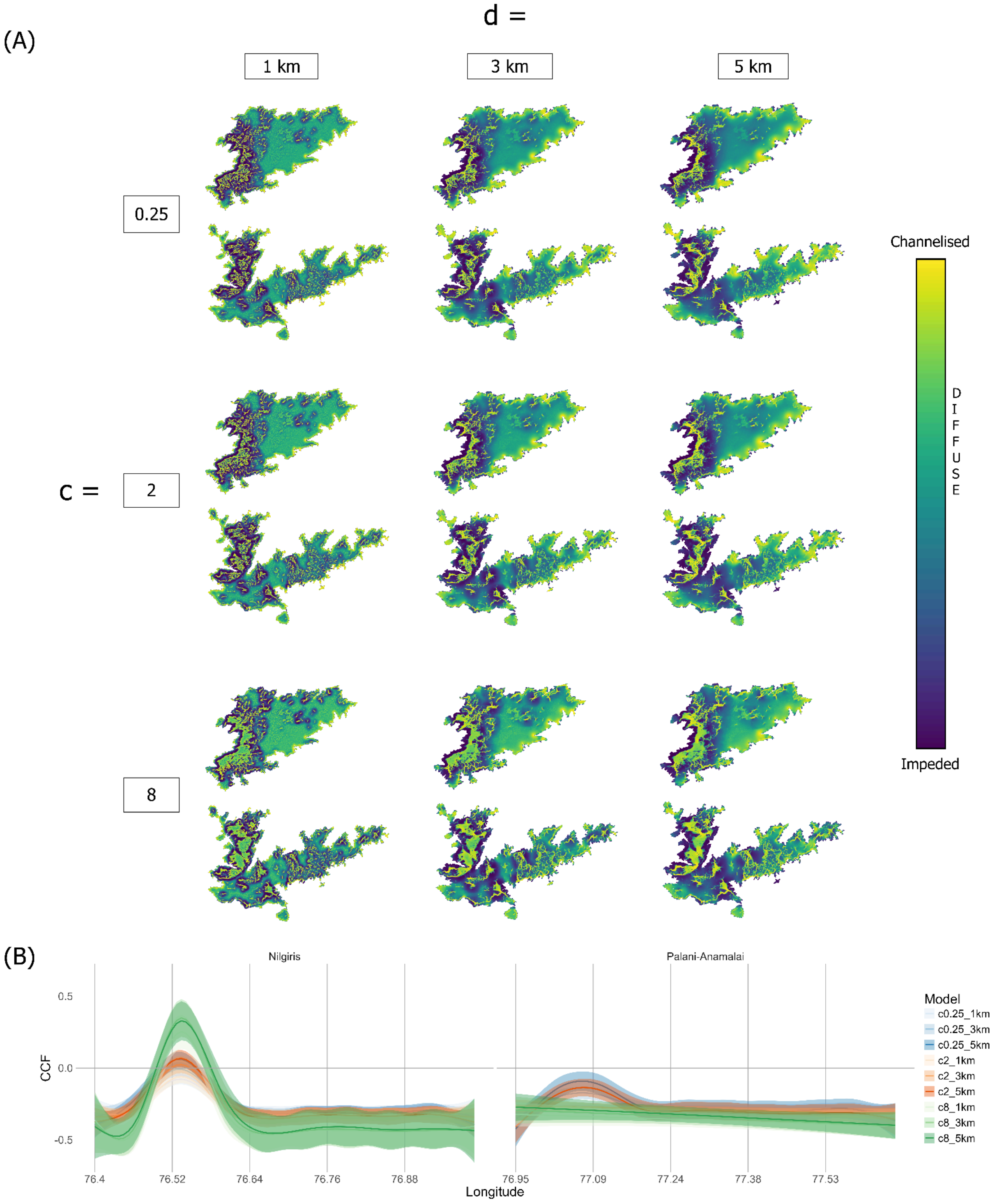


Fig. S9: Species - *Anthus nilghiriensis*. A) Present-day normalised current flow (NCF). Colour scale for visualisation is according to percentile bins for NCF values. NCF > 1 represents channelised current flow or potential movement corridors (yellow colours), NCF < 1 represents impeded current flow or potential barriers to movement (dark blue colours), and NCF ≈ 1 represents diffuse current flow (green-ish blue colours), i.e., neither impeded nor channelised. B) Present-day cumulative current flow (CCF) transformed to represent multiples of standard deviations away from the mean for each model (*c* value x dispersal distance), over a longitudinal gradient going from west to east on the x-axis for each corresponding sky island. Standard deviation stretched values for CCF are plotted with a 95% confidence interval after smoothing using a Generalised Additive Model (GAM). *c* determines the non-linearity of the habitat suitability to resistance transformation used to generate the resistance layer, and *d* represents the radius of the moving window in the connectivity model.

### S3. Change in movement suitability vs. landscape change

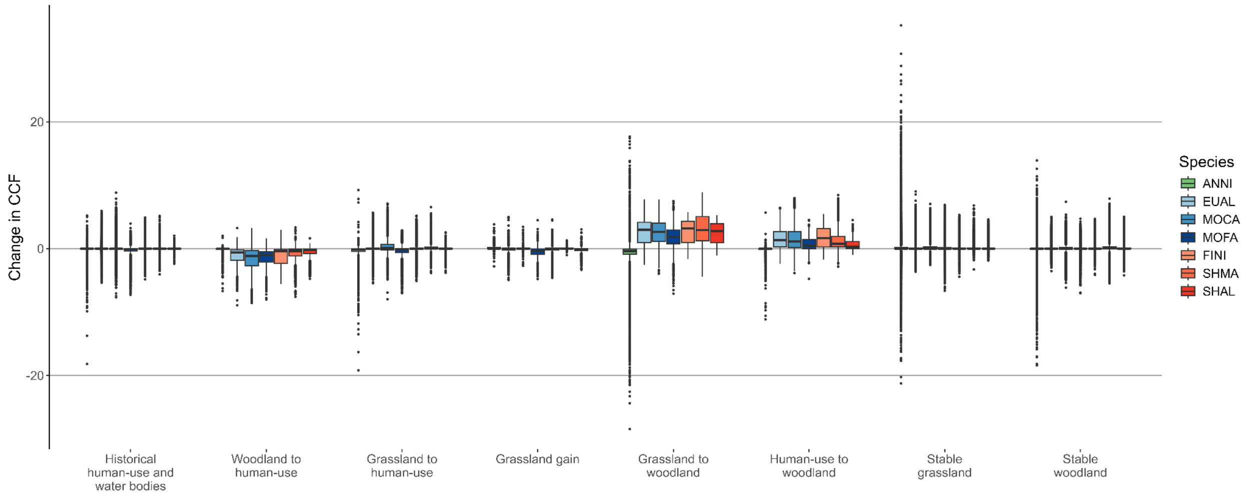


Fig. S10: Change in cumulative current flow (CCF) within 8 landscape change categories for different species, with outliers displayed. Values for change in CCF for each species have been transformed to represent multiples of standard deviations away from zero. Change in CCF > 0 represents an increase, Change in CCF < 0 represents a decrease, and Change in CCF ≈ 0 represents no change in current flow (movement suitability). Species key: ANNI - Anthus nilghiriensis (grassland specialist), EUAL - Eumyias albicaudatus (forest specialist), MOCA - Montecincla cachinnans (forest specialist), MOFA - Montecincla fairbanki (forest specialist), FINI - Ficedula nigrorufa (forest specialist), SHMA - Sholicola major (forest specialist), SHAL - Sholicola albiventris (forest specialist).

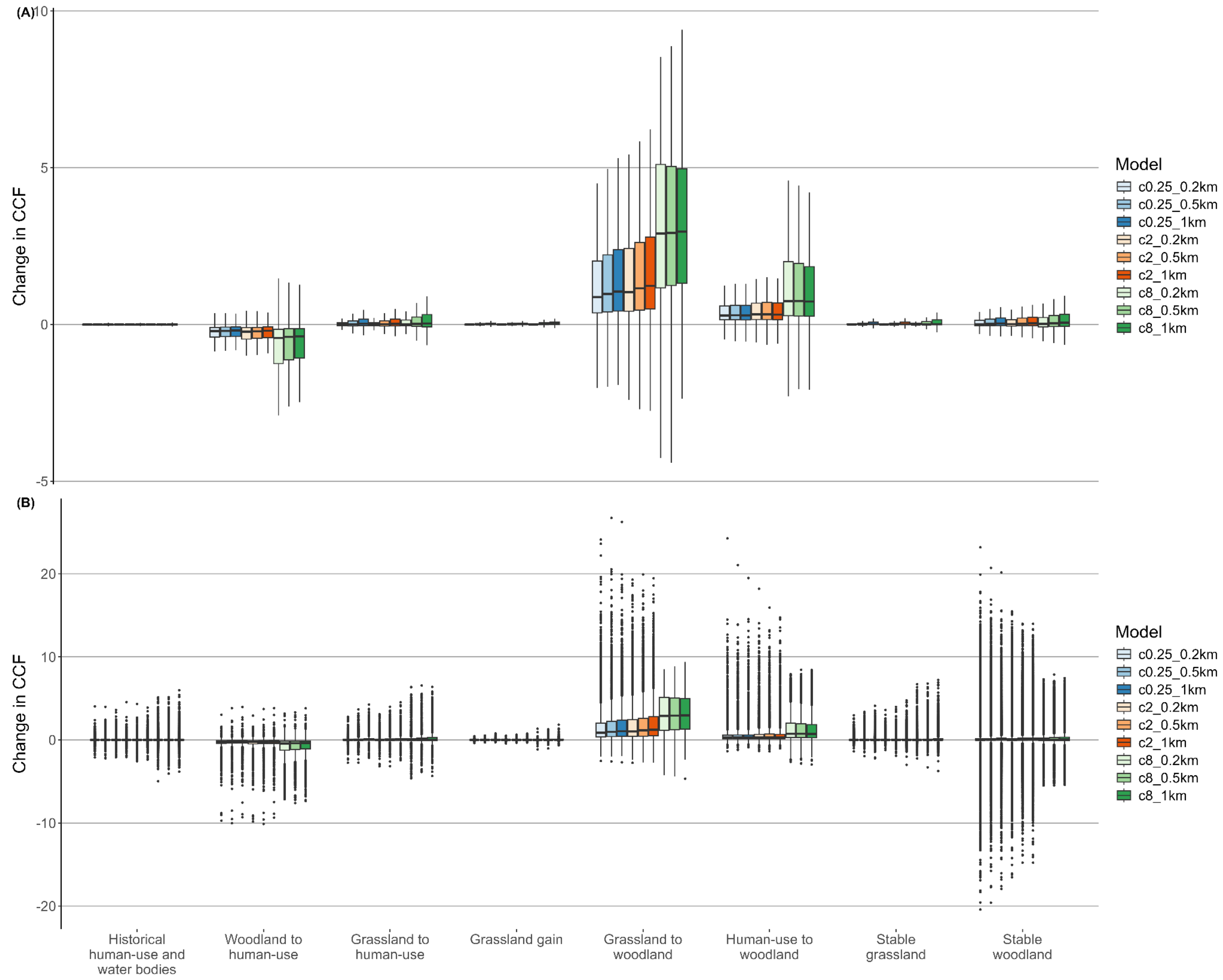


Fig. S11: Change in cumulative current flow (CCF) within 8 landscape change categories for each model (*c* value x dispersal distance) for *Sholicola major*, with A) outliers hidden, and B) outliers displayed. Values for change in CCF for each model have been transformed to represent multiples of standard deviations away from zero. Change in CCF > 0 represents an increase, Change in CCF < 0 represents a decrease, and Change in CCF ≈ 0 represents no change in current flow (movement suitability).

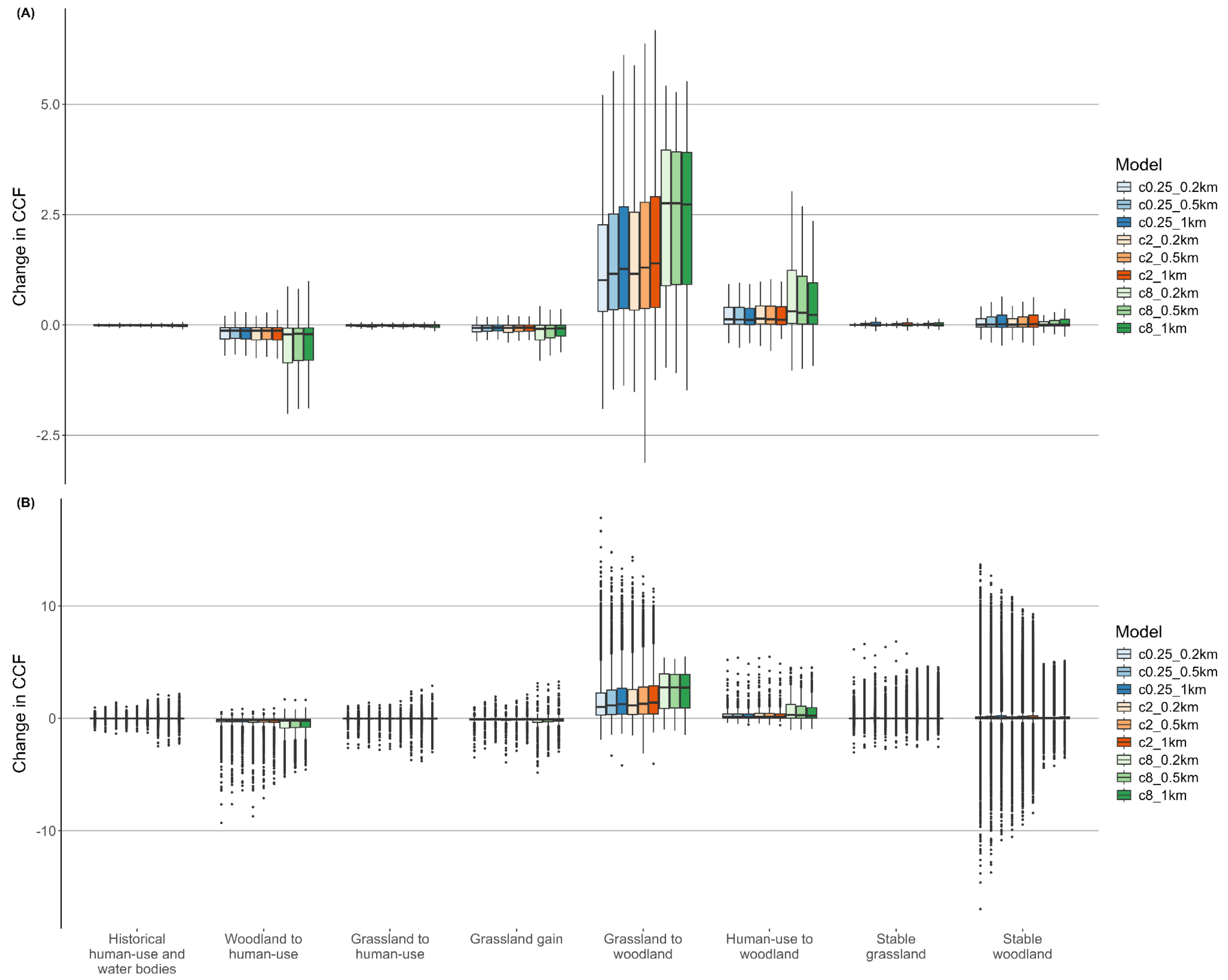


Fig. S12: Change in cumulative current flow (CCF) within 8 landscape change categories for each model (*c* value x dispersal distance) for *Sholicola albiventris*, with A) outliers hidden, and B) outliers displayed. Values for change in CCF for each model have been transformed to represent multiples of standard deviations away from zero. Change in CCF > 0 represents an increase, Change in CCF < 0 represents a decrease, and Change in CCF ≈ 0 represents no change in current flow (movement suitability).

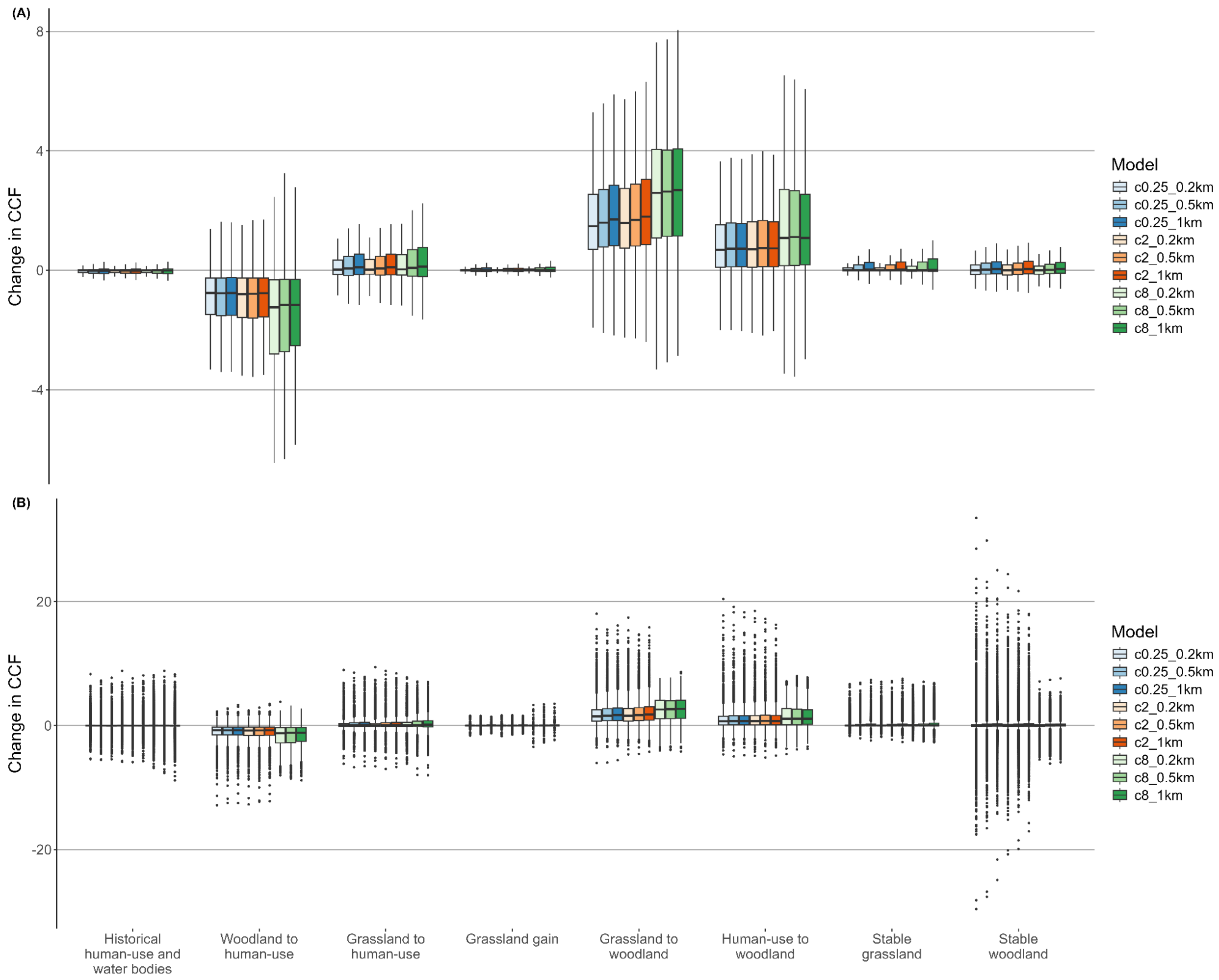


Fig. S13: Change in cumulative current flow (CCF) within 8 landscape change categories for each model (*c* value x dispersal distance) for *Montecincla cachinnans*, with A) outliers hidden, and B) outliers displayed. Values for change in CCF for each model have been transformed to represent multiples of standard deviations away from zero. Change in CCF > 0 represents an increase, Change in CCF < 0 represents a decrease, and Change in CCF ≈ 0 represents no change in current flow (movement suitability).

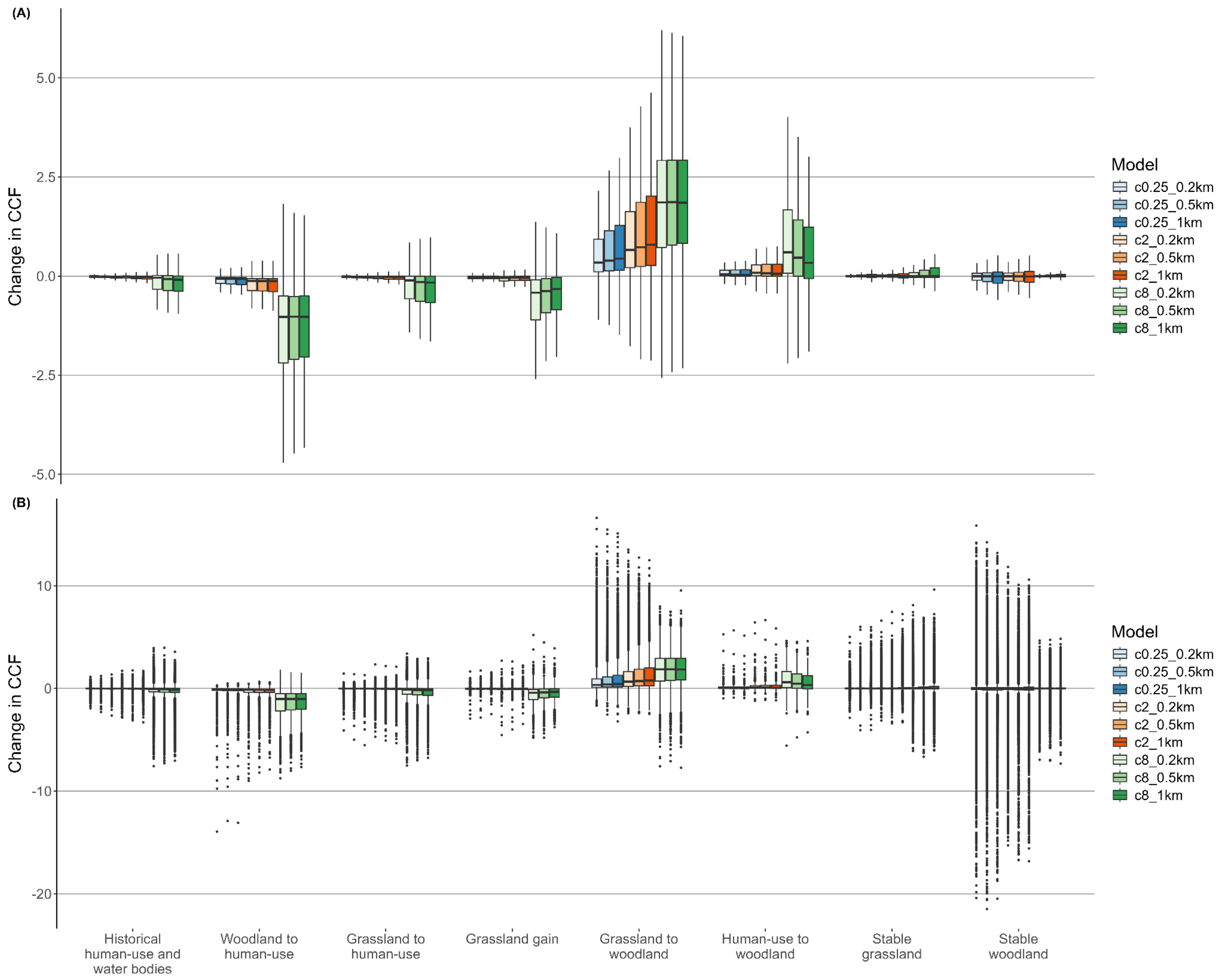


Fig. S14: Change in cumulative current flow (CCF) within 8 landscape change categories for each model (*c* value x dispersal distance) for *Montecincla fairbanki*, with A) outliers hidden, and B) outliers displayed. Values for change in CCF for each model have been transformed to represent multiples of standard deviations away from zero. Change in CCF > 0 represents an increase, Change in CCF < 0 represents a decrease, and Change in CCF ≈ 0 represents no change in current flow (movement suitability).

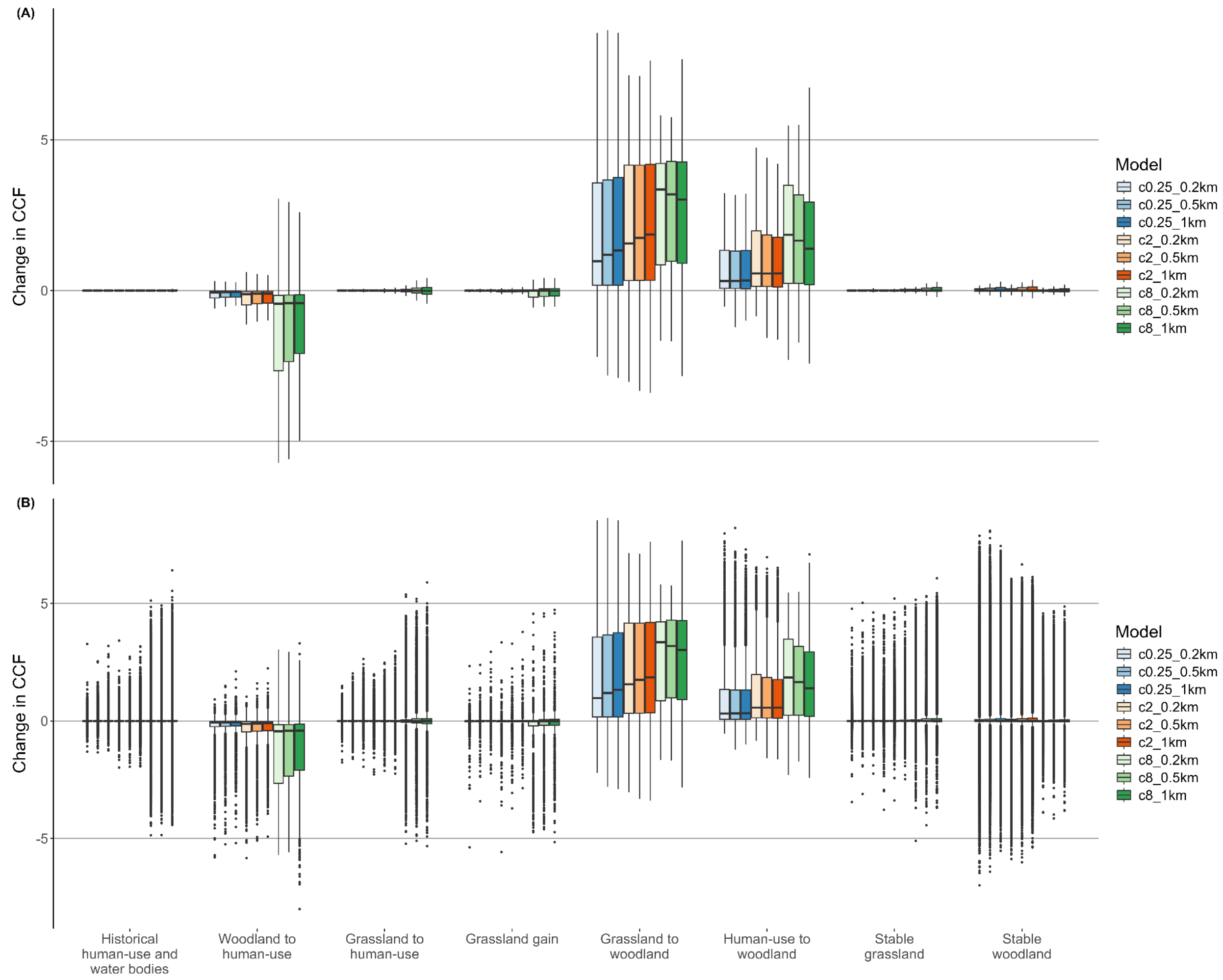


Fig. S15: Change in cumulative current flow (CCF) within 8 landscape change categories for each model (*c* value x dispersal distance) for *Ficedula nigrorufa*, with A) outliers hidden, and B) outliers displayed. Values for change in CCF for each model have been transformed to represent multiples of standard deviations away from zero. Change in CCF > 0 represents an increase, Change in CCF < 0 represents a decrease, and Change in CCF ≈ 0 represents no change in current flow (movement suitability).

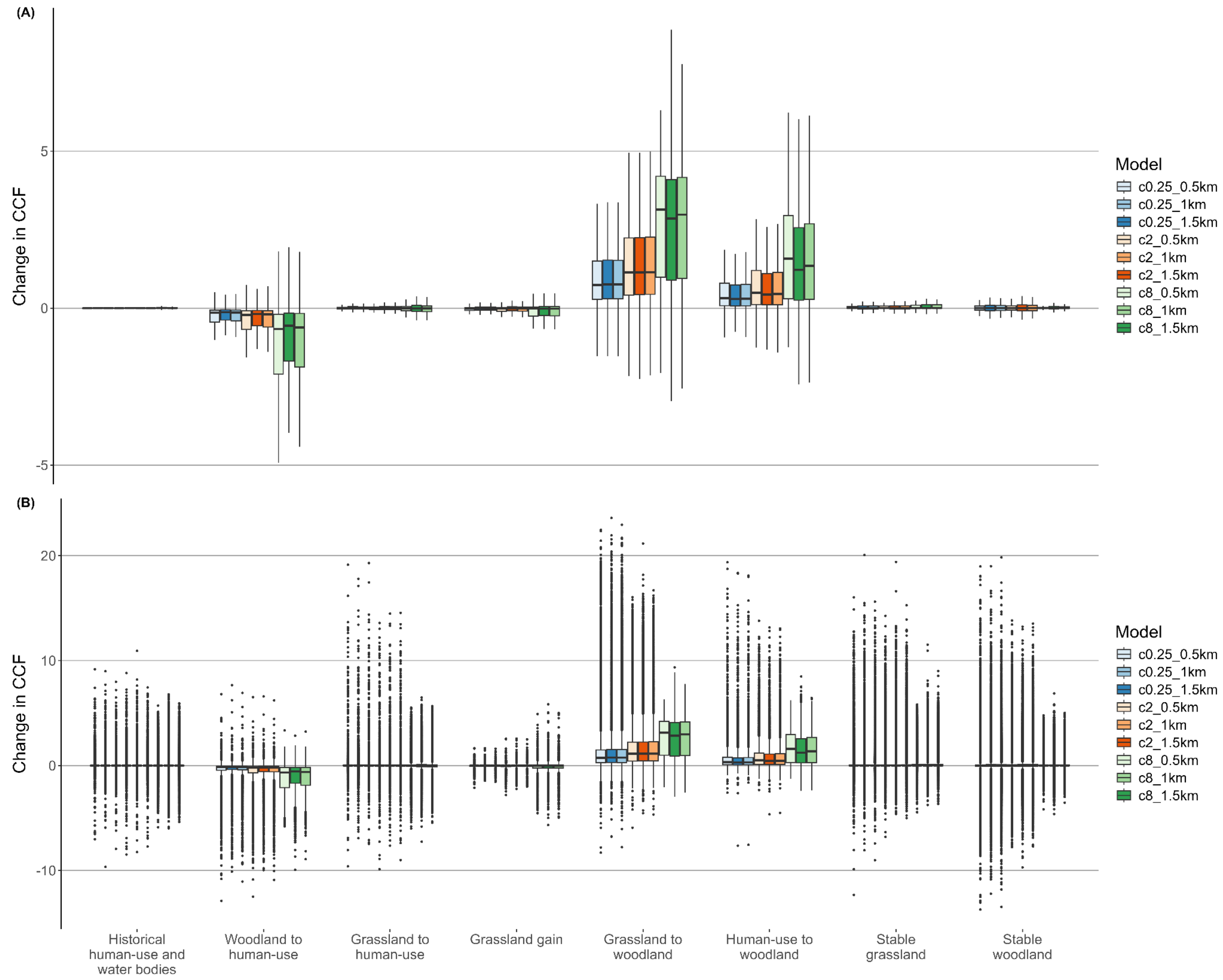


Fig. S16: Change in cumulative current flow (CCF) within 8 landscape change categories for each model (*c* value x dispersal distance) for *Eumyias albicaudatus*, with A) outliers hidden, and B) outliers displayed. Values for change in CCF for each model have been transformed to represent multiples of standard deviations away from zero. Change in CCF > 0 represents an increase, Change in CCF < 0 represents a decrease, and Change in CCF ≈ 0 represents no change in current flow (movement suitability).

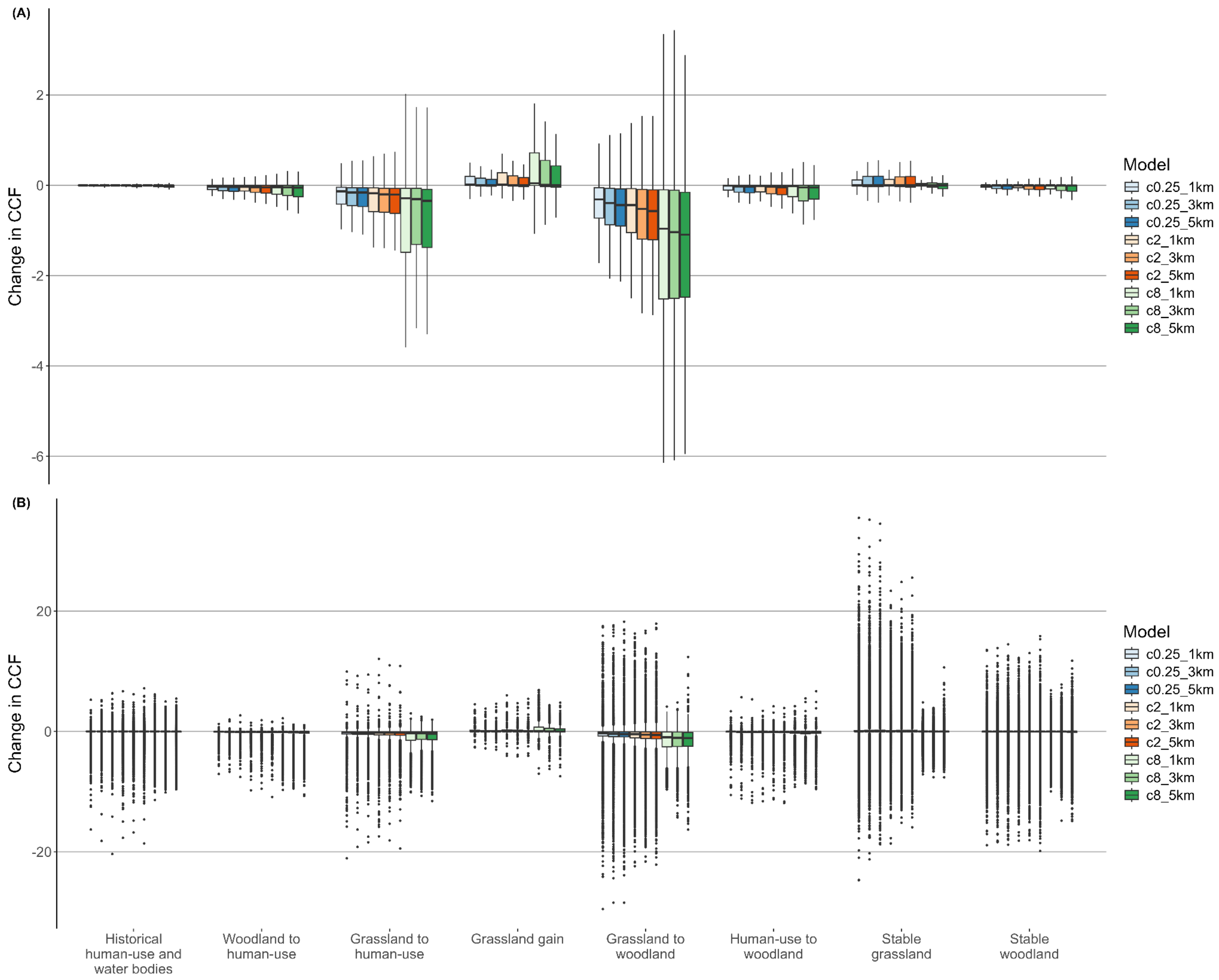


Fig. S17: Change in cumulative current flow (CCF) within 8 landscape change categories for each model (*c* value x dispersal distance) for *Anthus nilghiriensis*, with A) outliers hidden, and B) outliers displayed. Values for change in CCF for each model have been transformed to represent multiples of standard deviations away from zero. Change in CCF > 0 represents an increase, Change in CCF < 0 represents a decrease, and Change in CCF ≈ 0 represents no change in current flow (movement suitability).

### S4. Species-specific changes in spatial patterns of movement

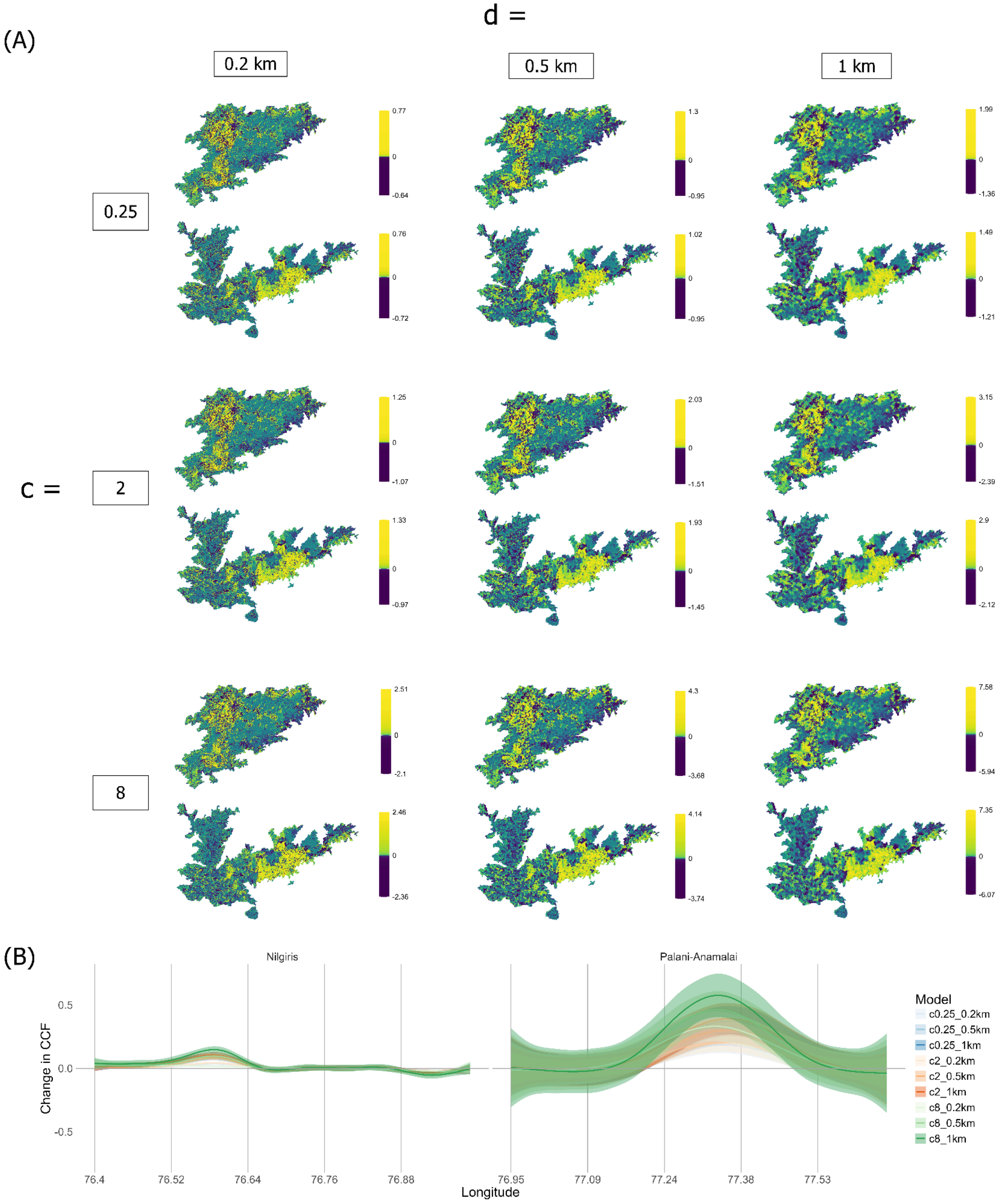


Fig. S18: Species - *Sholicola major* (in the Nilgiris) and *Sholicola albiventris* (in the Palani-Anamalais). A) Maps for change in CCF for each model. Colour scale for visualisation is according to percentile bins for change in CCF values. B) Change in CFF for each model over a longitudinal gradient, going from west to east on the x-axis for each corresponding Sky Island. Values for change in CCF for each model have been transformed to represent multiples of standard deviations away from zero. Change in CCF $> 0$ represents an increase, Change in CCF $< 0$ represents a decrease, and Change in CCF $\approx 0$ represents no change in current flow (movement suitability). Standard deviation stretched values for change in CCF are plotted with a 95% confidence interval after smoothing using a Generalised Additive Model (GAM). *c* determines the non-linearity of the habitat suitability to resistance transformation used to generate the resistance layer, and *d* represents the radius of the moving window in the connectivity model.

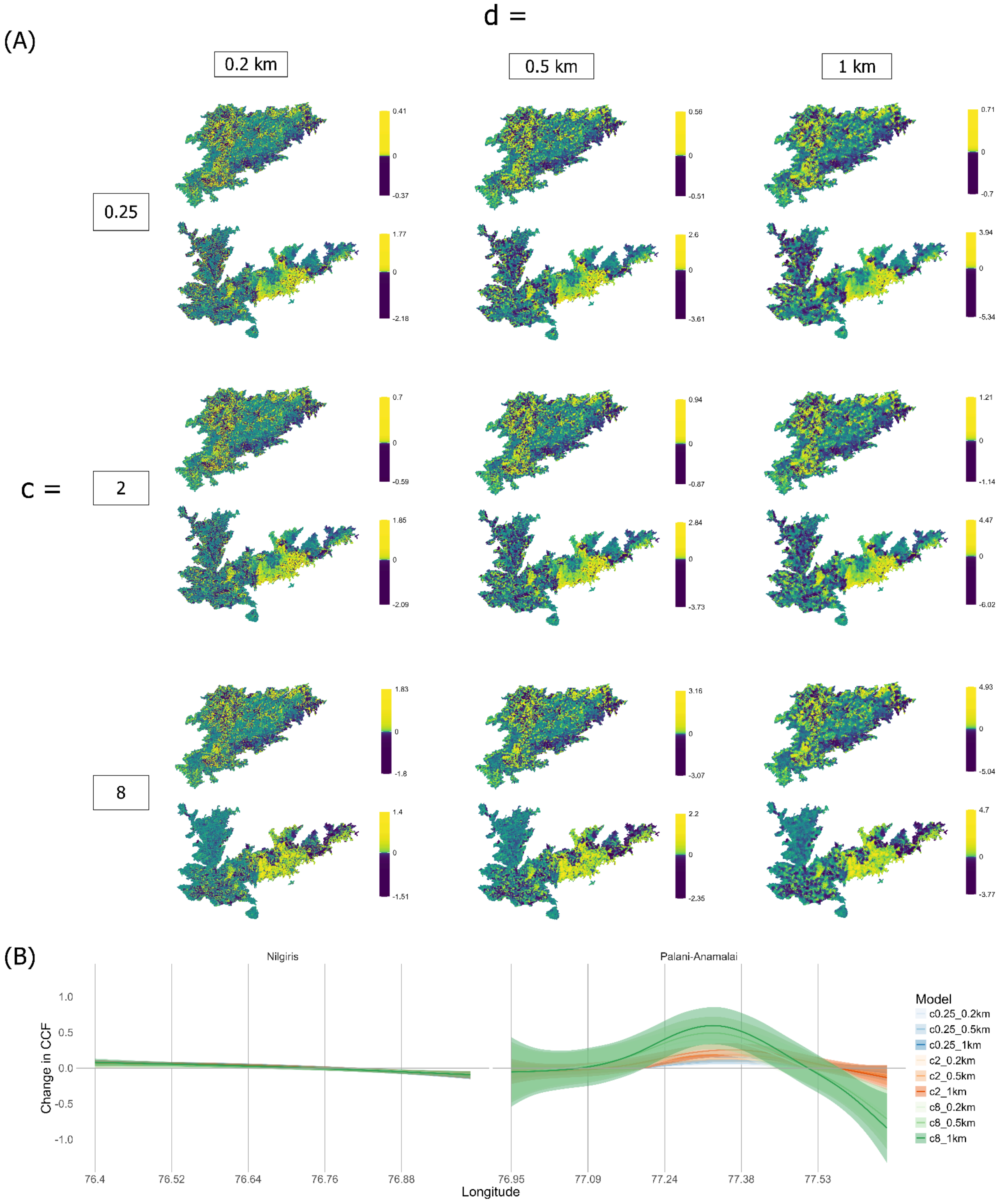


Fig. S19: Species - *Montecincla cachinnans* (in the Nilgiris) and *Montecincla fairbanki* (in the Palani-Anamalais). A) Maps for change in CCF for each model. Colour scale for visualisation is according to percentile bins for change in CCF values. B) Change in CFF for each model over a longitudinal gradient, going from west to east on the x-axis for each corresponding Sky Island. Values for change in CCF for each model have been transformed to represent multiples of standard deviations away from zero. Change in CCF > 0 represents an increase, Change in CCF < 0 represents a decrease, and Change in CCF ≈ 0 represents no change in current flow (movement suitability). Standard deviation stretched values for change in CCF are plotted with a 95% confidence interval after smoothing using a Generalised Additive Model (GAM). *c* determines the non-linearity of the habitat suitability to resistance transformation used to generate the resistance layer, and *d* represents the radius of the moving window in the connectivity model.

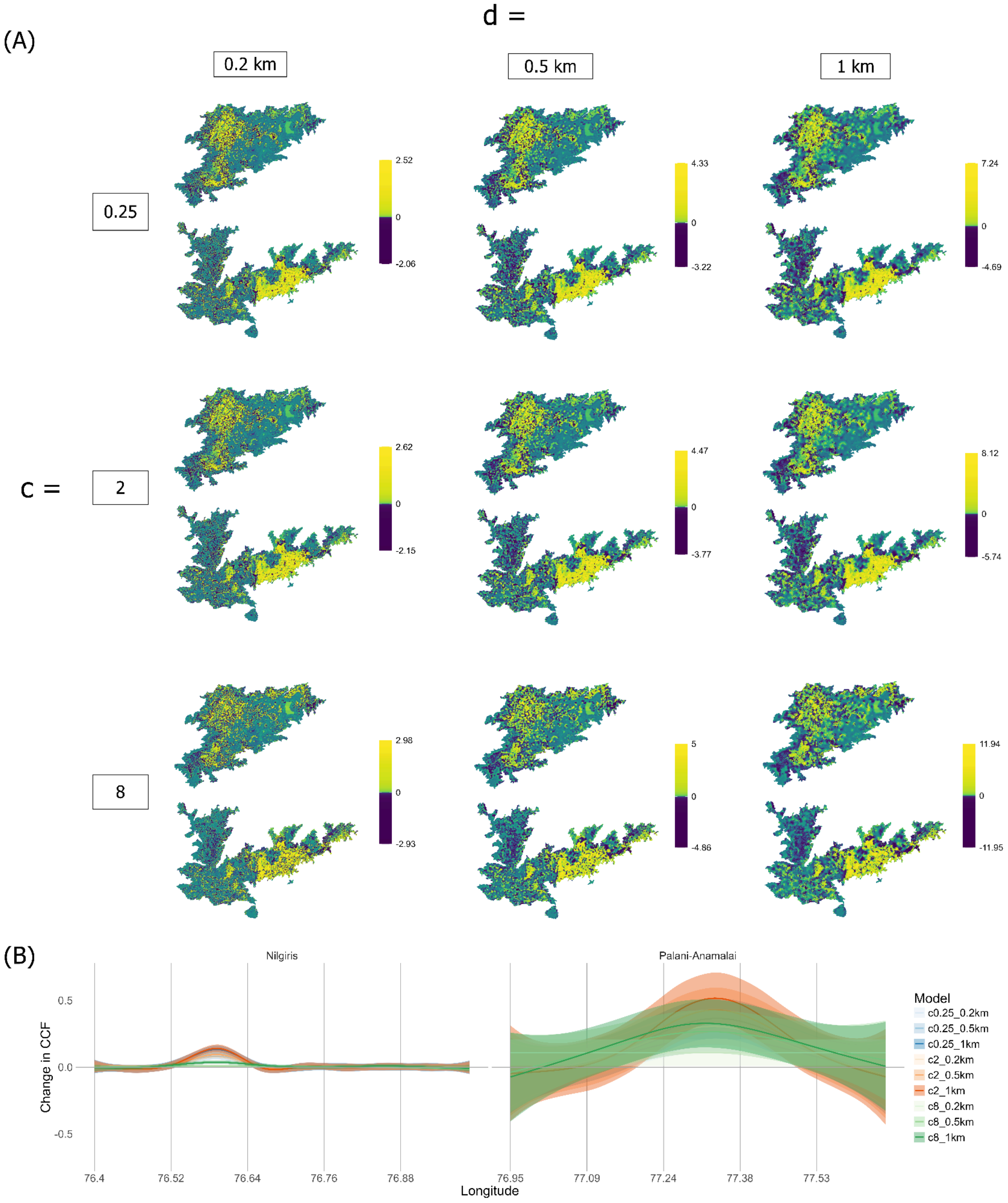


Fig. S20: Species - *Ficedula nigrorufa*. A) Maps for change in CCF for each model. Colour scale for visualisation is according to percentile bins for change in CCF values. B) Change in CFF for each model over a longitudinal gradient, going from west to east on the x-axis for each corresponding Sky Island. Values for change in CCF for each model have been transformed to represent multiples of standard deviations away from zero. Change in CCF > 0 represents an increase, Change in CCF < 0 represents a decrease, and Change in CCF ≈ 0 represents no change in current flow (movement suitability). Standard deviation stretched values for change in CCF are plotted with a 95% confidence interval after smoothing using a Generalised Additive Model (GAM). *c* determines the non-linearity of the habitat suitability to resistance transformation used to generate the resistance layer, and *d* represents the radius of the moving window in the connectivity model.

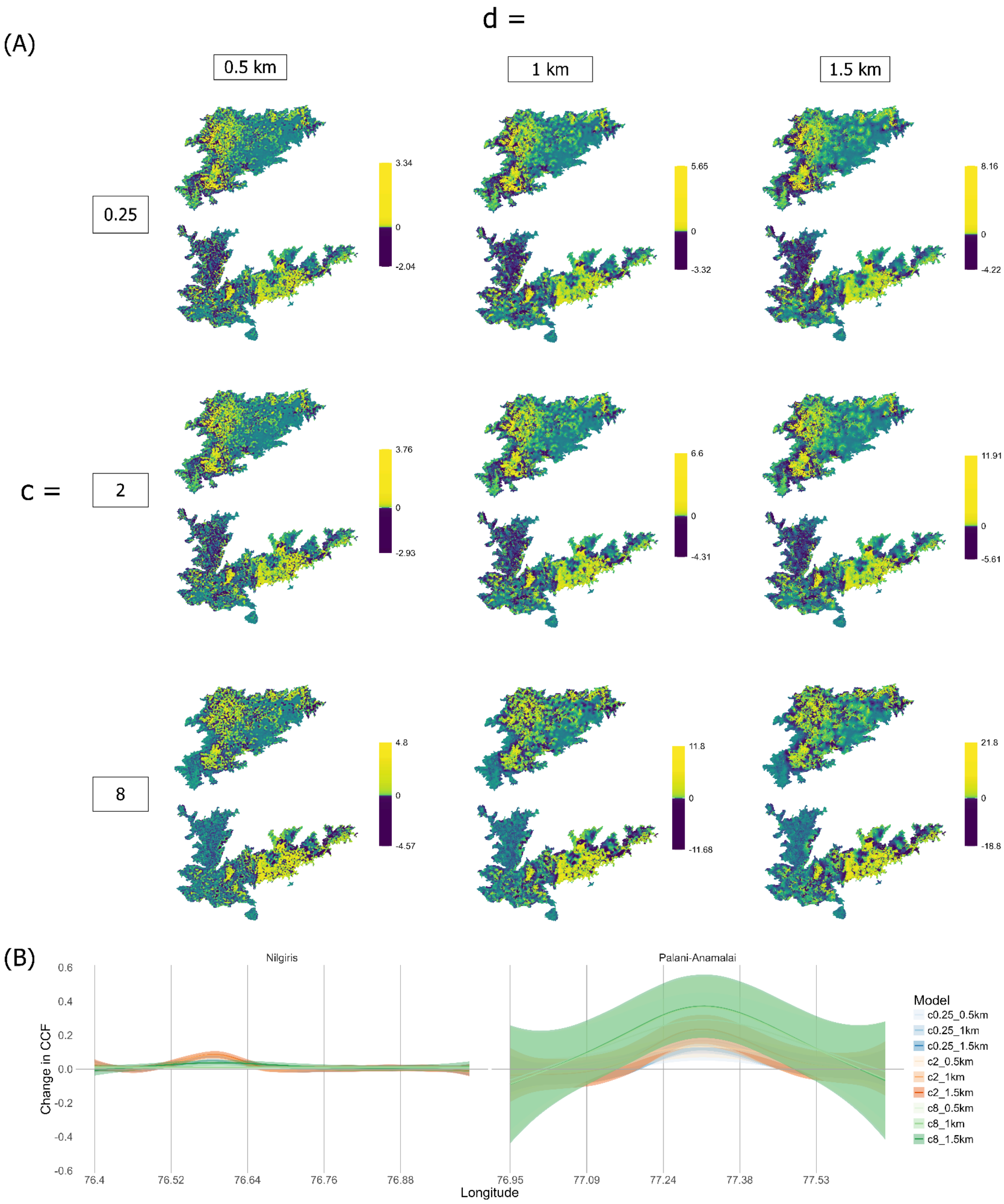


Fig. S21: Species - *Eumyias albicaudatus*. A) Maps for change in CCF for each model. Colour scale for visualisation is according to percentile bins for change in CCF values. B) Change in CFF for each model over a longitudinal gradient, going from west to east on the x-axis for each corresponding Sky Island. Values for change in CCF for each model have been transformed to represent multiples of standard deviations away from zero. Change in CCF > 0 represents an increase, Change in CCF < 0 represents a decrease, and Change in CCF ≈ 0 represents no change in current flow (movement suitability). Standard deviation stretched values for change in CCF are plotted with a 95% confidence interval after smoothing using a Generalised Additive Model (GAM). *c* determines the non-linearity of the habitat suitability to resistance transformation used to generate the resistance layer, and *d* represents the radius of the moving window in the connectivity model.

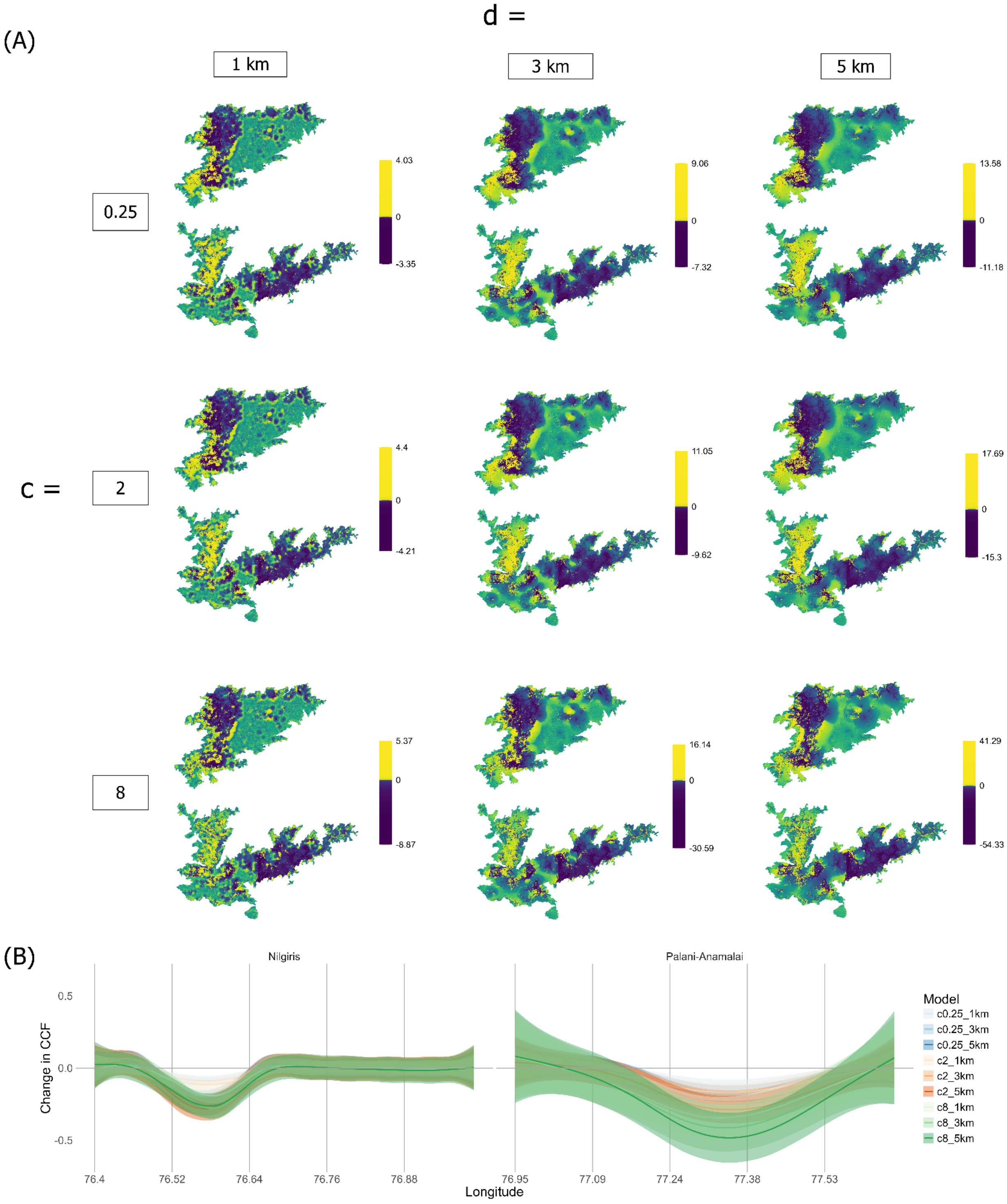


Fig. S22: Species - *Anthus nilghiriensis*. A) Maps for change in CCF for each model. Colour scale for visualisation is according to percentile bins for change in CCF values. B) Change in CFF for each model over a longitudinal gradient, going from west to east on the x-axis for each corresponding Sky Island. Values for change in CCF for each model have been transformed to represent multiples of standard deviations away from zero. Change in CCF > 0 represents an increase, Change in CCF < 0 represents a decrease, and Change in CCF ≈ 0 represents no change in current flow (movement suitability). Standard deviation stretched values for change in CCF are plotted with a 95% confidence interval after smoothing using a Generalised Additive Model (GAM). *c* determines the non-linearity of the habitat suitability to resistance transformation used to generate the resistance layer, and *d* represents the radius of the moving window in the connectivity model.
